# Endogenous viral element-derived piRNAs are not required for production of ping-pong-dependent piRNAs from Diaphorina citri densovirus

**DOI:** 10.1101/2020.05.20.105924

**Authors:** Jared C. Nigg, Yen-Wen Kuo, Bryce W. Falk

## Abstract

Partial integrations of DNA and non-retroviral RNA virus genomes, termed endogenous viral elements (EVEs), are abundant in arthropod genomes and often produce PIWI-interacting RNAs (piRNAs) speculated to target cognate viruses through the ping-pong cycle, a post-transcriptional RNA silencing mechanism. Here we describe a Diaphorina citri densovirus (DcDV)-derived EVE in the genome of *Diaphorina citri*. We found that this EVE gives rise to DcDV-specific primary piRNAs and is unevenly distributed among *D. citri* populations. Unexpectedly, we found that DcDV is targeted by ping-pong-dependent viral piRNAs (vpiRNAs) in *D. citri* lacking the DcDV-derived EVE, while four naturally infecting RNA viruses of *D. citri* are not targeted by vpiRNAs. Furthermore, a recombinant Cricket paralysis virus containing a portion of the DcDV genome corresponding to the DcDV-derived EVE was not targeted by vpiRNAs during infection in *D. citri* harboring the EVE. These results represent the first report of ping-pong-dependent vpiRNAs outside of mosquitoes.

## Introduction

The Piwi-interacting RNA (piRNA) pathway is a small RNA (sRNA) guided gene silencing mechanism responsible for repressing transposable elements (TEs) in animals and emerging evidence supports a role for this pathway in antiviral responses in some mosquito species and mosquito-derived cell lines (1–5). In *Drosophila melanogaster*, biogenesis of primary piRNAs begins with transcription of single-stranded piRNA precursor transcripts from discrete genomic regions called piRNA clusters (6). TE sequences integrated into piRNA clusters can become transcribed as part of a piRNA precursor transcript and these precursor transcripts are processed into primary piRNAs that direct transcriptional and post-transcriptional silencing of TEs by association with the Piwi-family Argonaute proteins Piwi and Aubergine, respectively (1). During the ping-pong cycle, cleavage of sense TE RNA directed by Aubergine-bound antisense primary piRNAs triggers production of sense secondary piRNAs from cleaved TE RNA and, in turn, sense secondary piRNAs direct cleavage of antisense piRNA precursor transcripts via association with another Piwi-family Argonaute protein, Argonaute 3 (Ago3), to specifically amplify the response against active TEs (6). piRNAs are distinguished from other sRNAs by their size (24-32 nt), nucleotide biases (uridine as the first nucleotide for primary piRNAs and adenine as the tenth nucleotide for secondary piRNAs, known as the 1U and 10A bias), and association with a Piwi-family Argonaute protein (1). Additionally, complementary piRNAs produced by the ping-pong cycle have 5’ ends separated by exactly 10 nt, known as the ping-pong signature (1).

The presence of ping-pong-dependent virus-derived piRNAs (vpiRNAs) during infection with several RNA viruses in *Aedes* and *Culex* mosquitoes and cell lines has been reported (2–5,7–12) and reduced expression of piRNA pathway components leads to increased replication of Semliki Forest virus, Bunyamwera virus, Cache Valley virus, and Rift Valley Fever virus in *Aedes aegypti-*derived Aag2 cells (2–5). Despite the prevalence of virus-derived piRNAs and evidence supporting their antiviral role, little is known regarding their biogenesis or antiviral function *in vivo*. *Ae. aegypti* expresses an expanded group of Piwi-family Argonaute proteins containing eight proteins (Piwi1-7 and Ago3) rather than the three proteins seen in *D. melanogaster* (Piwi, Aub, and Ago3) (13). In Aag2 cells, Piwi5 and Ago3 are the primary Piwi-family Argonaute proteins required for biogenesis of vpiRNAs upon infection with Sindbis virus (9). Importantly, induction of the ping-pong cycle in mosquitoes is thought to involve recognition of exogenous viral RNA, triggering biogenesis of primary vpiRNAs directly from viral RNA without the need for primary piRNAs derived from endogenous loci (9, 11). This is in contrast to the canonical ping-pong cycle, which produces primary piRNAs only from endogenous transcripts (6). Thus, mosquitoes possess a unique piRNA pathway that produces vpiRNAs by virtue of novel functionality that has so far not been found in other groups of organisms. Indeed, the piRNA pathway was not found to play an antiviral role in *D. melanogaster* and no evidence for an antiviral piRNA pathway, or even the presence of ping-pong-dependent vpiRNAs, has been reported outside of the mosquito lineage (14).

Arthropod genomes contain an abundance of sequences derived from DNA viruses and non-retroviral RNA viruses (15–20). These sequences, termed endogenous viral elements (EVEs), are often enriched within piRNA clusters and are known to serve as sources of primary piRNAs in a wide variety of arthropod species (15–17). These observations have led to speculation that EVE-derived primary piRNAs may play an antiviral role by targeting cognate viruses in a manner analogous to piRNA-mediated repression of TEs via the ping-pong cycle (15–17). Importantly, such a role for EVE-derived piRNAs could theoretically be mediated by the canonical ping-pong cycle and could thus expand the antiviral potential of the piRNA pathway beyond mosquitoes. In support of this idea, Whitfield et al. found that a single EVE-derived piRNA maps to the genome of Phasi Charoen-like virus (PCLV) in Aag2 cells, which are persistently infected with PCLV, and that a vpiRNA is produced from the complementary strand (16). Moreover, the authors found that knockdown of Piwi4 led to a ∼2 fold increase in the abundance of PCLV RNA in Aag2 cells (16). While these results are intriguing, it is difficult to assess how the EVE-derived piRNA affects PCLV RNA abundance in the context of the Piwi4 knockdown, as only global sRNA mapping data to the entire PCLV nucleocapsid-encoding segment are presented, the abundance of the EVE-derived piRNA and its potential PCLV-derived ping-pong partner were not discussed, and Piwi4 knockdown has been shown to reduce both siRNA and piRNA production (16, 21). More recently, Tassetto et al. found that Piwi4 is required for vpiRNA maturation in Aag2 cells and binds specifically to piRNAs derived from EVEs (21). Furthermore, they found that insertion of EVE sequences into the 3’ UTR of Sindbis virus reduced viral replication in a Piwi4-dependent manner in Aag2 cells (21). These results raise the possibility that EVE-derived piRNAs may target cognate viruses in these cells through an interaction with Piwi4, however, detailed sRNA profiles and more thorough examinations of the interplay between virus-specific piRNAs derived from endogenous and exogenous sources are needed to establish the link between EVE-derived piRNAs and an antiviral response.

We previously identified the EVEs present within the genomes of 48 arthropod species (15). Among the EVEs that we estimated to be similar enough to their corresponding exogenous viruses at the nucleotide level such that piRNAs derived from them could potentially target viral RNA during infection, Diaphorina citri densovirus (DcDV) and a DcDV-derived EVE located within a piRNA cluster on *Diaphorina citri* genomic scaffold 2850 stood out as an EVE-virus pair sharing exceptionally high nucleotide identity over a relatively long region (15). *D. citri*, also known as the Asian citrus psyllid, is a hemipteran insect pest and serves as a vector of *Candidatus* Liberibacter species that cause Huanglongbing, or citrus greening disease, in all types of commercially cultivated citrus throughout the world (22). This disease represents the greatest threat to the citrus industry worldwide and has devastated many citrus growing regions, such as in the US state of Florida, where citrus production has decreased by 74% since the first report of citrus greening disease in the state in 2005 (23). Control strategies for the disease are limited and primarily involve removal of infected trees and management of *D. citri* populations with chemical insecticides (24). The continued spread of citrus greening disease and the development of insecticide resistance in field populations of *D. citri* highlight the need for new management approaches (24). Biological control strategies relying on infectious agents have proven to be effective methods for controlling insect pests and the use of parasites and viruses have been proposed as possible vector control approaches for *D. citri* (25–28). Current understanding of immune processes in *D. citri* is limited and studies in this field are needed to facilitate the development of effective biological control strategies. In particular, antiviral mechanisms in *D. citri* have not been experimentally evaluated. The presence of a highly similar virus-EVE pair, a lack of novel Piwi-family of Argonaute proteins like those observed in mosquitoes, and the need to study antiviral mechanisms in *D. citri* make this species an excellent and relevant model to study the interactions between EVE-derived piRNAs and corresponding viruses.

Here we further characterized the DcDV-derived EVE present within the *D. citri* genome. We found that piRNAs are produced from this EVE in multiple tissue types and that this EVE is conserved among some geographically distinct populations of *D. citri*, but is absent from other populations. By analyzing sRNA profiles in *D. citri* insects infected with DcDV, we found that this virus is targeted by ping-pong-dependent vpiRNAs independent of DcDV-derived EVEs and endogenous DcDV-specific piRNAs. Additionally, analysis of sRNA profiles during infection with other viruses suggests that RNA viruses are not a target of vpiRNAs in *D. citri*.

## Results

### A DcDV-derived EVE with high nucleotide identity to DcDV is conserved in some populations of D. citri, but absent in others

We previously identified a 621 bp DcDV-derived EVE located within a piRNA cluster on *D. citri* genomic scaffold 2850 by comparing deduced virus protein sequences to deduced *D. citri* genome-encoded protein sequences using BLASTx (15). This EVE was 85.5% identical to the corresponding region of the DcDV genome at the deduced amino acid level and resided just downstream from another 624 bp EVE derived from a different portion of the DcDV genome. To characterize these EVEs at the nucleotide level, we aligned the nucleotide sequence of the DcDV genome to the region of the *D. citri* genome harboring these DcDV-derived EVEs. We found that this region of the *D. citri* genome contains sequences corresponding to the DcDV inverted terminal repeats (ITRs) and to two regions of the non-structural protein (NS) gene cassette together spanning 1,396 bp. For simplicity we refer to these regions as endogenous ITR (EITR) and endogenous NS (ENS) (Fig. 1a). Notably, the portion of ENS spanning nucleotides 7,523 to 8,175 within genomic scaffold 2850 shares 86% nucleotide identity with the corresponding region of the DcDV genome.

**Fig. 1.**
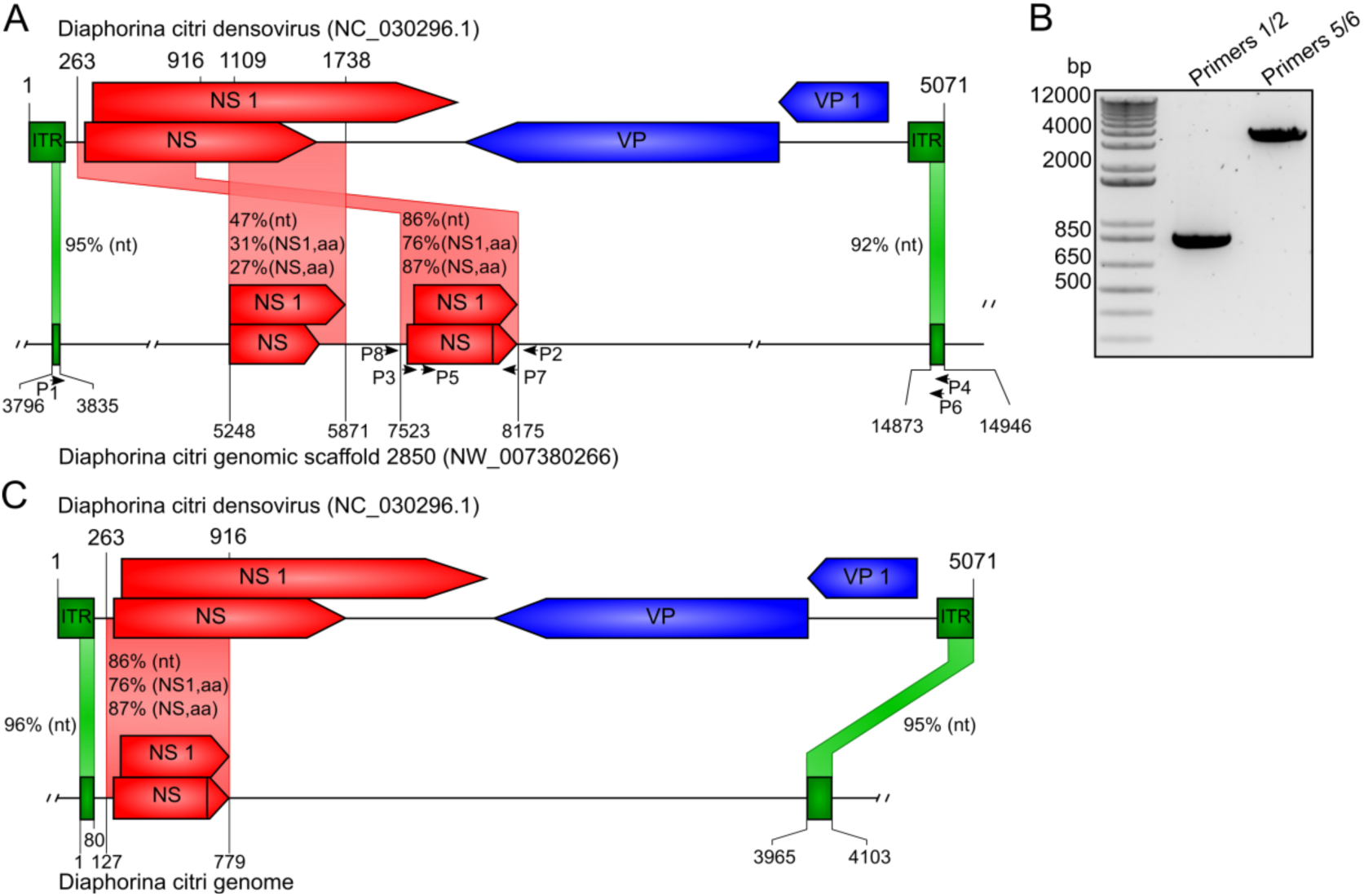
A DcDV-like EVE is present in the *D. citri* genome. (A) Organization of DcDV-derived EVEs identified in *D. citri* genomic scaffold 2850 by BLASTx followed by manual sequence alignment. The DcDV genome organization is shown on top with the corresponding region of scaffold 2850 on the bottom. Numbers above and below the sequence depictions represent nucleotide positions. Vertical lines inside shaded boxes represent stop codons. The percent nucleotide or deduced amino acid (aa) identity between EVEs and corresponding viral genomic regions/proteins is given. Annealing positions of PCR primers are shown with arrows. (B) Confirmation of EVE presence by PCR using primers shown in panel A. The product produced with primers 5/6 was produced by nested PCR using as template a 1:1000 dilution of a PCR product produced with primers 3/4. (C) The correct organization of the region of the *D. citri* genome containing ENS and EITR, as confirmed by sequencing of the PCR products shown in panel B.

Because our EVE prediction shown in Fig. 1a was based only on BLAST results and a previous *D. citri* genome assembly, we wanted to confirm the presence, sequence, and organization of the DcDV-derived EVEs. To confirm the presence of these EVEs, we designed PCR primers to amplify this region from *D. citri* genomic DNA (Fig. 1a). We obtained PCR products of unexpected size (based on the available sequence of scaffold 2850 and the organization depicted in Fig. 1a) for both primer sets (Fig. 1b). Sequencing of these PCR products revealed that the portion of ENS spanning nucleotides 5,248 to 5,871 within genomic scaffold 2850 was not present and the distance between the ITR-like fragments was 3,885 bp rather than 11,038 bp. Additionally, EITR was 210 bp instead of 114 bp. The portion of ENS corresponding to the region between DcDV nucleotides 263-916 was present and closely matched the sequence observed by BLAST. Figure 1c shows the correct organization of this region of the DcDV genome, as confirmed by sequencing of the PCR products shown in Fig. 1b. This organization and sequence is supported by a more recent draft of the *D. citri* genome assembled using PacBio long reads (Diaci2.0, ftp://ftp.citrusgreening.org/genomes/Diaphorina_citri/assembly/DIACI_v2.0/).

Our PCR reactions were performed using DNA extracted from *D. citri* insects collected from a colony located at the University of California Davis Contained Research Facility (CRF) that was originally started using insects collected in the US state of California (this colony is designated CRF-CA). The *D. citri* reference genome used for BLAST searches and nucleotide alignments was produced by sequencing DNA from insects collected in the US state of Florida (29). Haplotype network analysis of global *D. citri* populations based on the mitochondrial cytochrome oxidase subunit I gene indicates the existence of 44 *D. citri* haplotypes belonging to two distinct lineages, denoted lineages A and B (30–32). The invasion history of *D. citri* out of Southern Asia has resulted in segregation of the two lineages, such that only lineage B is found in North America, while lineage A predominates in Southeast Asia, Africa, and South America (30). Besides CRF-CA, three other *D. citri* colonies are maintained at the CRF and these were started using *D. citri* insects collected in Taiwan, Uruguay, and the US state of Hawaii (designated CRF-TW, CRF-Uru, and CRF-HI, respectively). To determine whether ENS is conserved among geographically distinct *D. citri* populations, we performed PCR using primers flanking ENS and DNA extracted from insects collected from CRF-CA, CRF-TW, CRF-Uru, and CRF-HI. We also included DNA extracted from field collected *D. citri* insects from Pakistan, Brazil, and the US states of Arizona and Florida. We obtained nearly identical PCR products from insects from CRF-CA, CRF-HI, Pakistan, Arizona, and Florida, but no PCR products from insects from CRF-TW, CRF-Uru, or Brazil (Fig. 2a & S1). Based on the known distribution of the two *D. citri* lineages, these results suggest that ENS is absent in *D. citri* from lineage A.

**Fig 2.**
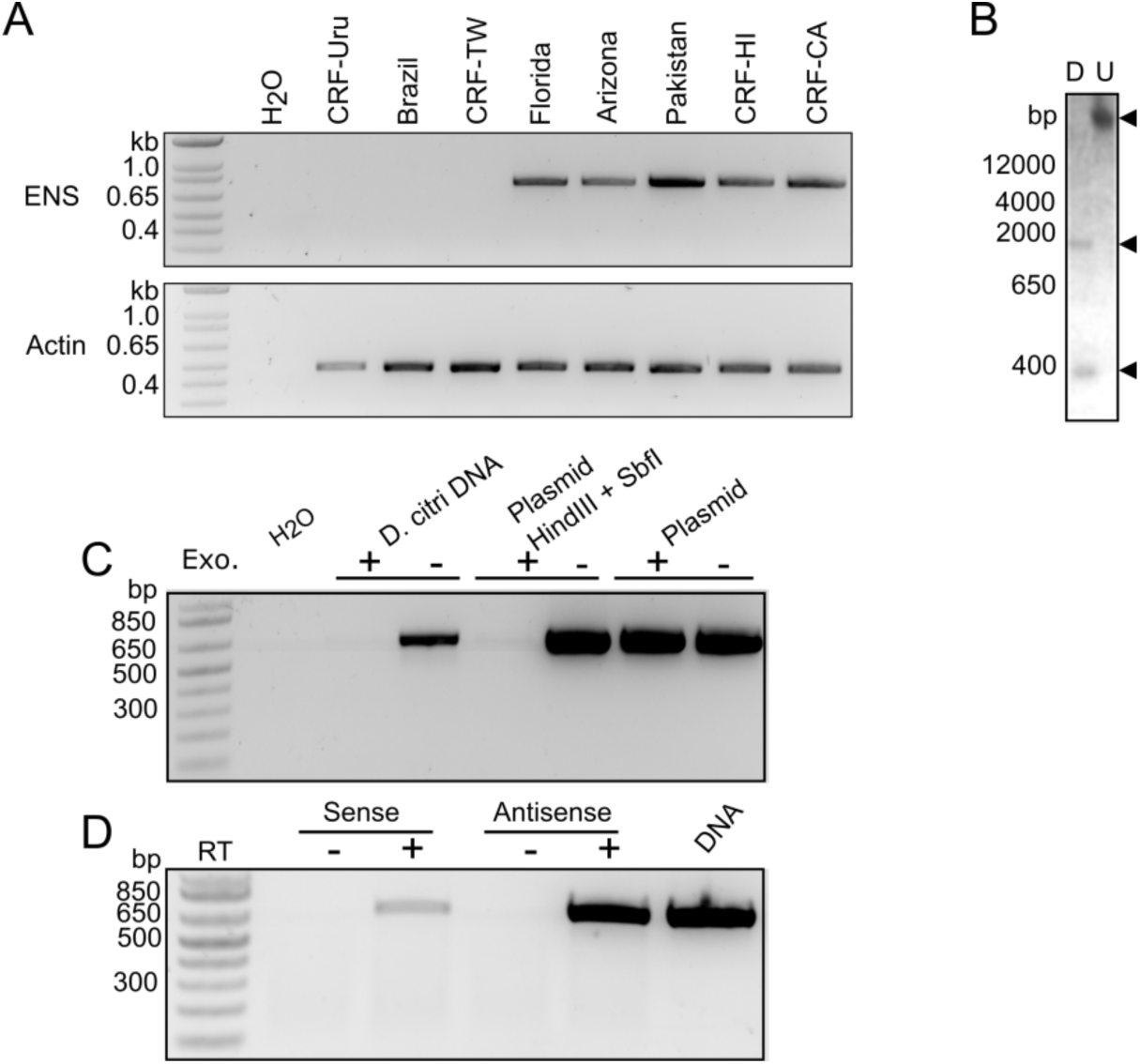
ENS is a transcribed EVE that is unevenly distributed among distinct *D. citri* populations. (A) Upper: PCR products produced using primers flanking ENS (primers 2 and 8). Lower: PCR products produced using primers specific to *D. citri* actin (primers 9 and 10). (B) Southern blot of *D. citri* genomic DNA using an RNA probe based on the sequence of ENS; U = undigested, D = digested with PstI and HindIII. Upper arrow denotes ENS in undigested genomic DNA. Lower arrows denote cleavage products. (C) PCR products produced using primers 3 and 7 and the indicated DNA samples. “Plasmid” is a plasmid containing the full ENS sequence and “Plasmid HindIII + SbfI” is the same plasmid digested with HindIII and SbfI. DNA was left intact or digested with an exonuclease (Exo.) prior to PCR. (D) Primers 3 or 7 were used to generate cDNA from antisense or sense transcripts, respectively. cDNAs were used as templates for PCR using primers 3 and 7. DNA or cDNA prepared without reverse transcriptase (RT) served as controls.

Sequences derived from some insect infecting viruses are maintained as circular episomal molecules that produce virus-specific sRNAs (12, 33). Given the heterogenous distribution of ENS among *D. citri* populations, we examined whether ENS is maintained episomally rather than representing a true EVE. When we performed a Southern blot with DNA extracted from CRF-CA *D. citri* using an RNA probe based on the sequence of ENS we observed a single DNA species at high molecular weight (>12,000 bp) from undigested DNA and two species of lower molecular weight in DNA digested with PstI and HindIII, indicating that ENS resides in high molecular weight genomic DNA rather than low molecular weight episomal DNA (Fig. 2b). The observation of two DNA species in the digested sample does not represent a second DcDV-like EVE because ENS contains a PstI clevage site. To confirm that ENS is a true EVE, we treated *D. citri* genomic DNA with an exonuclease to remove genomic DNA, but not circular episomal DNA, and performed PCR with ENS-specific primers using exonuclease-treated DNA as template. We obtained a PCR product from non-digested DNA, but not from digested DNA (Fig. 2c). Together, these results indicate the ENS is indeed integrated into the *D. citri* genome. Finally, strand specific RT-PCR indicates that ENS is bidirectionally transcribed, although the majority of transcripts are antisense to the corresponding DcDV transcript (Fig. 2d).

### ENS and EITR give rise to DcDV-specific primary piRNAs

To evaluate whether ENS and EITR give rise to virus-specific primary piRNAs as described for other EVEs, we mapped sRNAs from CRF-CA *D. citri* not infected with DcDV to the DcDV genome. We found that sRNAs purified from CRF-CA *D. citri* mapped to EITR and to the negative strand within the portion of the DcDV genome corresponding to ENS (i.e. antisense to DcDV transcripts), but not to other regions (Fig. 3a). The size of these sRNAs was characteristic of piRNAs and they possessed a strong 1U bias (Fig. 3b & c). Similar results were obtained for other *D. citri* populations that are not infected with DcDV and for which sRNA datasets are available (Fig. S2a & b). As expected based on the lack of ENS in *D. citri* insects collected in Brazil or from CRF-Uru, DcDV-specific piRNAs were not observed in these insects (Fig. S2c & d). We note that the DcDV ITRs are comprised of a 210 nt hairpin present on both ends of the DcDV genome. Thus, it is not possible to determine whether the sRNAs mapping to EITR are specific to the positive or the negative strand. For this reason, because of the relatively small size of EITR compared to ENS, and because EITR does not correspond to a transcribed region of the DcDV genome (34), we chose to focus our analysis on piRNAs derived from ENS.

**Fig. 3.**
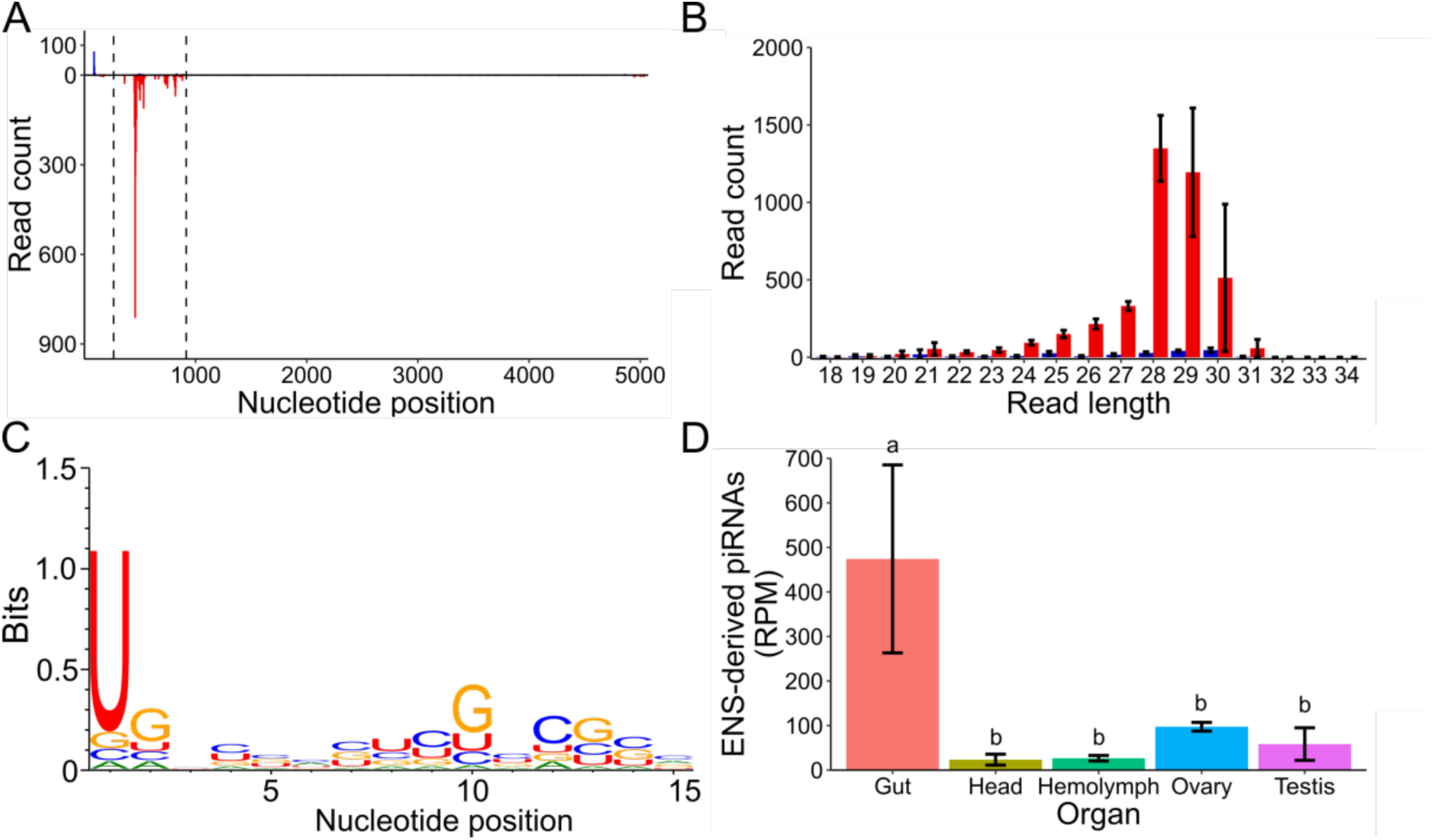
DcDV-specific piRNAs are produced from ENS in *D. citri* from CRF-CA. (A) Positions of all sRNAs from CRF-CA *D. citri* mapped to the DcDV genome. Dashed lines indicate the region of the DcDV genome corresponding to ENS. sRNAs are shown as mapped to the genomic strand containing the coding sequence for the NS proteins. Red = antisense sRNAs, blue = sense sRNAs. Reads counts are an average of three independent libraries. (B) Length distribution of sRNAs shown in (A). Red = antisense, blue = sense. Read counts are an average of three independent libraries. Error bars indicate standard deviation. (C) Sequence logo of sRNAs shown in (A). Data represents three pooled libraries. (D) Abundance of 27-32 nt sRNAs from CRF-CA *D. citri* mapping to ENS in the indicated tissue types. Read counts for each library were normalized using the read counts per million mapped reads method (RPM)(59). Normalized read counts represent the average of three independent libraries. Error bars indicate standard deviation. Average RPM values for each tissue were compared by one-way ANOVA and Turkey’s honestly significant difference post-hoc test. Significance is indicated by lowercase letters and tissues sharing a letter do not have significantly different RPM values (p < 0.05).

The expression of piRNAs is known to display tissue specificity and piRNA expression patterns could have important consequences for the ability of EVE-derived piRNAs to target cognate viruses. To determine the tissue specificity of piRNAs produced from ENS and to determine the tissues in which the ping-pong cycle is active in *D. citri*, we sequenced sRNAs purified from dissected *D. citri* guts, heads, ovaries, testis, and hemolymph using insects collected from CRF-CA. We found that while DcDV-specific piRNAs derived from ENS were present in all tissues analyzed, their expression was significantly higher in *D. citri* guts than in any other tissue (Fig. 3d). The ping-pong cycle is restricted to germline tissues in *D. melanogaster* (1), however a comprehensive analysis of somatic sRNAs mapping to TEs genome wide in 20 arthropod species in combination with ancestral state reconstruction indicates that somatic ping-pong amplification is widespread throughout arthropods despite having been independently lost in some species, including *D. melanogaster* (35). To determine the tissues in which the ping-pong cycle is active in *D. citri* we mapped sRNAs from CRF-CA *D. citri* guts, heads, hemolymph, ovaries, and testis to all TEs identified within the *D. citri* genome and analyzed the mapped 27-32 nt sRNAs for the presence of ping-pong signatures. We found evidence for ping-pong amplification of TE-derived piRNAs in all tissues examined (Fig. S3).

### DcDV is targeted by ping-pong-dependent piRNAs in the absence of a DcDV-derived EVE

We previously found that *D. citri* from CRF-CA are resistant to infection with DcDV (34). In contrast, the virus is maintained as a persistent, maternally transmitted infection in *D. citri* from CRF-TW (34). Thus, to understand the sRNA-based response to DcDV infection in *D. citri*, we sequenced sRNAs from DcDV-infected *D. citri* from CRF-TW. These results revealed a major population of 21 nt sRNAs, indicative of an siRNA-based response (Fig. 4a). Unexpectedly, we also observed a smaller peak within the piRNA size range centered at 29 nt and we obtained 99.5% coverage of transcribed regions of the DcDV genome by mapping only 27-32 nt sRNAs (Fig. 4a & b). We found that complementary 27-32 nt sRNAs mapping to opposite strands throughout the DcDV genome possessed 5’ ends separated by 10 nt more often than expected by chance, an indication of ping-pong amplification (Z-score = 4.05±0.08)(Fig. 4c). Moreover, we detected the 1U and 10A biases typical of ping-pong amplification in 27-32 nt sRNAs mapping antisense and sense to the canonical DcDV transcripts, respectively (Fig. 4d & e).

**Fig. 4.**
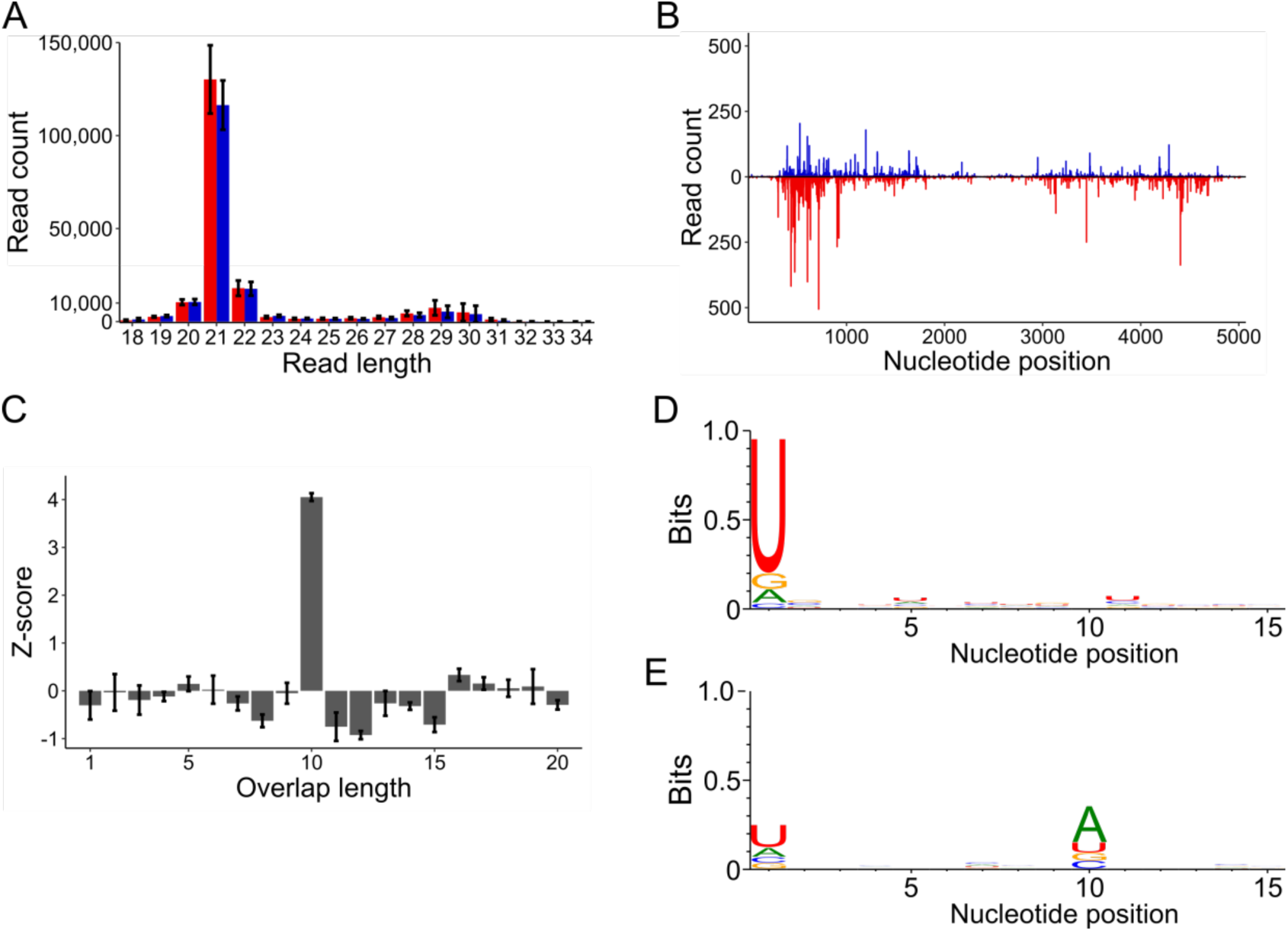
DcDV is targeted by ping-pong-dependent vpiRNAs in DcDV-infected *D. citri* insects from CRF-TW. (A) Length distribution of sRNAs mapping to the DcDV genome in DcDV-infected *D. citri* insects from CRF-TW. To account for the bidirectional transcription strategy of DcDV, sRNA mapping polarity was assigned from mapping location based on the start and stop positions of the canonical DcDV transcripts. Red = antisense sRNAs, blue = sense sRNAs. Read counts are an average of three independent libraries. Error bars indicate standard deviation. (B) Positions of 27-32 nt sRNAs shown in (A). Red = antisense sRNAs, blue = sense sRNAs. Read counts are an average of three independent libraries. (C) Z-scores for the indicated overlap distances between the 5’ ends of complementary 27-32 nt sRNAs shown in (A). Z-scores are an average of three independent libraries. Error bars indicate standard deviation. (D & E) Sequence logos for the 27-32 nt sRNAs shown in (A). Sequence logos for antisense sRNAs (D) or sense sRNAs (E) are shown. Data represents three pooled libraries.

We found that ENS is not present in the genome of CRF-TW *D. citri* (Fig. 2a), thus our observation of ping-pong-dependent DcDV-derived piRNAs in these insects suggests that DcDV is targeted by piRNAs independent of ENS. However, because this result was obtained by performing PCR using primers flanking ENS, it is possible that ENS is present in CRF-TW *D. citri,* but resides in different genomic context than seen in other *D. citri* populations. ENS is not identical to the corresponding region of the DcDV genome at the nucleotide level (Fig. 1c). Thus, some piRNAs derived from ENS do not perfectly map to DcDV, providing a means with which to distinguish some ENS-derived piRNAs from DcDV-derived piRNAs. To rule out the possibility that ENS is present in the genome of CRF-TW *D. citri*, we mapped 27-32 nt sRNAs from CRF-TW *D. citri* to the DcDV genome without allowing any mismatches. When the unmapped reads from this analysis were mapped to ENS without allowing any mismatches, no reads mapped. In contrast, we obtained an average of 55.8% coverage of the ENS sequence when the same analysis was performed using sRNAs from CRF-CA *D. citri* (data not shown). This result indicates that ENS is indeed not present in the genome of CRF-TW *D. citri* and that the targeting of DcDV by ping-pong-dependent vpiRNAs in these insects is independent of ENS-derived piRNAs.

We cannot exclude the possibility that the genome of CRF-TW *D. citri* harbors a different piRNA-producing DcDV-derived EVE. Because DcDV is maternally transmitted to 100% of the progeny of CRF-TW females (34), it is not possible to determine the repertoire of endogenous piRNAs in these insects in the absence of DcDV infection. Thus, we analyzed the sRNAs present in *D. citri* from CRF-Uru, as these insects lack ENS and are not infected with DcDV (Fig. 2a). We found that no DcDV-specific piRNAs are produced in uninfected CRF-Uru *D. citri*, indicating that these insects do not harbor a piRNA-producing DcDV-derived EVE (Fig. S2d). The absence of DcDV-specific piRNAs was not due to a lack of piRNAs in general, as we detected abundant ping-pong-dependent TE-derived piRNAs in these insects (Fig. S4). In contrast to *D. citri* from CRF-CA which are resistant to DcDV infection, we found that *D. citri* from CRF-Uru were susceptible to DcDV infection by both oral acquisition and intrathoracic injection (Fig. S5a & b)(34). Moreover, the progeny of CRF-Uru *D. citri* infected with DcDV were also infected with the virus (Fig. S5c). Analysis of sRNAs from the progeny of CRF-Uru *D. citri* that had been infected with DcDV by intrathoracic injection revealed the presence of ping-pong-dependent DcDV-derived piRNAs (ping-pong Z-score = 2.55) (Fig. S6). Together, these results demonstrate that DcDV is targeted by ping-pong-dependent vpiRNAs in two populations of *D. citri* independent of EVE-derived piRNAs.

### Other viruses of D. citri are not targeted by piRNAs

Besides DcDV, there are five other viruses known to infect *D. citri*: Diaphorina citri reovirus (double-stranded RNA), Diaphorina citri picrona-like virus (plus strand single-stranded RNA; + ssRNA), Diaphorina citri bunyavirus (minus strand single-stranded RNA), Diaphorina citri-associated c virus (+ ssRNA), and Diaphorina citri flavi-like virus (+ ssRNA)(28,36–38). To determine whether piRNAs are part of a general response to viruses in *D. citri*, we mapped sRNAs from publicly available sRNA libraries known to be derived from insects infected with one or more of each of these viruses to the corresponding viral genomes (with the exception of Diaphorina citri-associated c virus for which no such sRNA library exists) (28, 36). As expected, we detected a prominent peak at 21 nt for each virus, indicating a siRNA-based response (Fig. S7). While virus-derived sRNAs within the piRNA size range were present during infection with all four viruses, there were no peaks above background levels within the piRNA size range and 27-32 nt reads lacked signatures typical of primary or ping-pong-dependent piRNAs (Fig. S7). These results indicate that viruses in general are not targeted by piRNAs in *D. citri* and suggest that the targeting of DcDV by the piRNA pathway in *D. citri* may be due to differences in the infection cycles between RNA and DNA viruses.

### A recombinant CrPV-based reporter virus carrying DcDV sequence is not targeted by piRNAs in D. citri that produce primary piRNAs from a DcDV-derived EVE

As has been observed for RNA viruses in mosquitoes, our results suggest that DcDV is targeted by ping-pong-dependent vpiRNAs due to the *de novo* production of vpiRNAs from exogenous DcDV RNA. If EVE-derived piRNAs were to prime the ping-pong cycle in this context, it would be difficult to distinguish priming driven by EVE-derived piRNAs from priming driven by virus-derived vpiRNAs. Because our results suggest that RNA viruses are not targeted by vpiRNAs in *D. citri*, we sought to construct a recombinant RNA virus harboring DcDV-derived EVE sequence in order to study potential priming of the ping-pong cycle by EVE-derived piRNAs without the background of vpiRNAs produced directly from viral RNA. For this purpose, we used Cricket paralysis virus (CrPV), a dicistrovirus with a + ssRNA genome that was originally isolated from field crickets (39). Due to the broad experimental host range of CrPV, including *D. citri* (E Matsumura, personal communication, October 2018), and the availability of an infectious clone, CrPV is often used to study antiviral mechanisms in insects (40, 41).

To directly test the hypothesis that EVE-derived primary piRNAs can prime ping-pong amplification during infection with viruses sharing complementary sequence, we inserted 57 nt from the DcDV genome (from a region corresponding to ENS) into the CrPV genome between the 1A and 2B coding sequences flanked by duplicate copies of the 1A cleavage site (Fig. 5a, Fig. S8). The recombinant DcDV sequence was inserted into the CrPV genome such that ENS-derived primary piRNAs mapping antisense to DcDV transcripts would map to the negative strand of the recombinant CrPV-DcDV genome. Thus, in both cases, ENS-derived primary piRNAs are complementary to the viral coding sequence and in analogy to the mechanisms of ping-pong amplification in the context of TEs, should be theoretically capable of directing cleavage of viral mRNAs. This recombinant virus is designated CrPV-DcDV. While the 57 nt inserted region is relatively small in the context of the ∼9 kb CrPV genome, piRNA target sites corresponding to single piRNAs are known to be sufficient for induction of ping-pong amplification and a single piRNA produced from a PCLV-derived EVE was proposed to mediate a piRNA-based antiviral response in Aag2 cells (16,42–44). Additional analysis of the sRNA mapping data from DcDV-uninfected CRF-CA *D. citri* shown in Figure 3a indicates that this 57 nt region gives rise to an average of 171.7 27-32 nt sRNAs made up of an average of 30.7 unique sequences in the three sRNA libraries analyzed (data not shown).

**Fig. 5.**
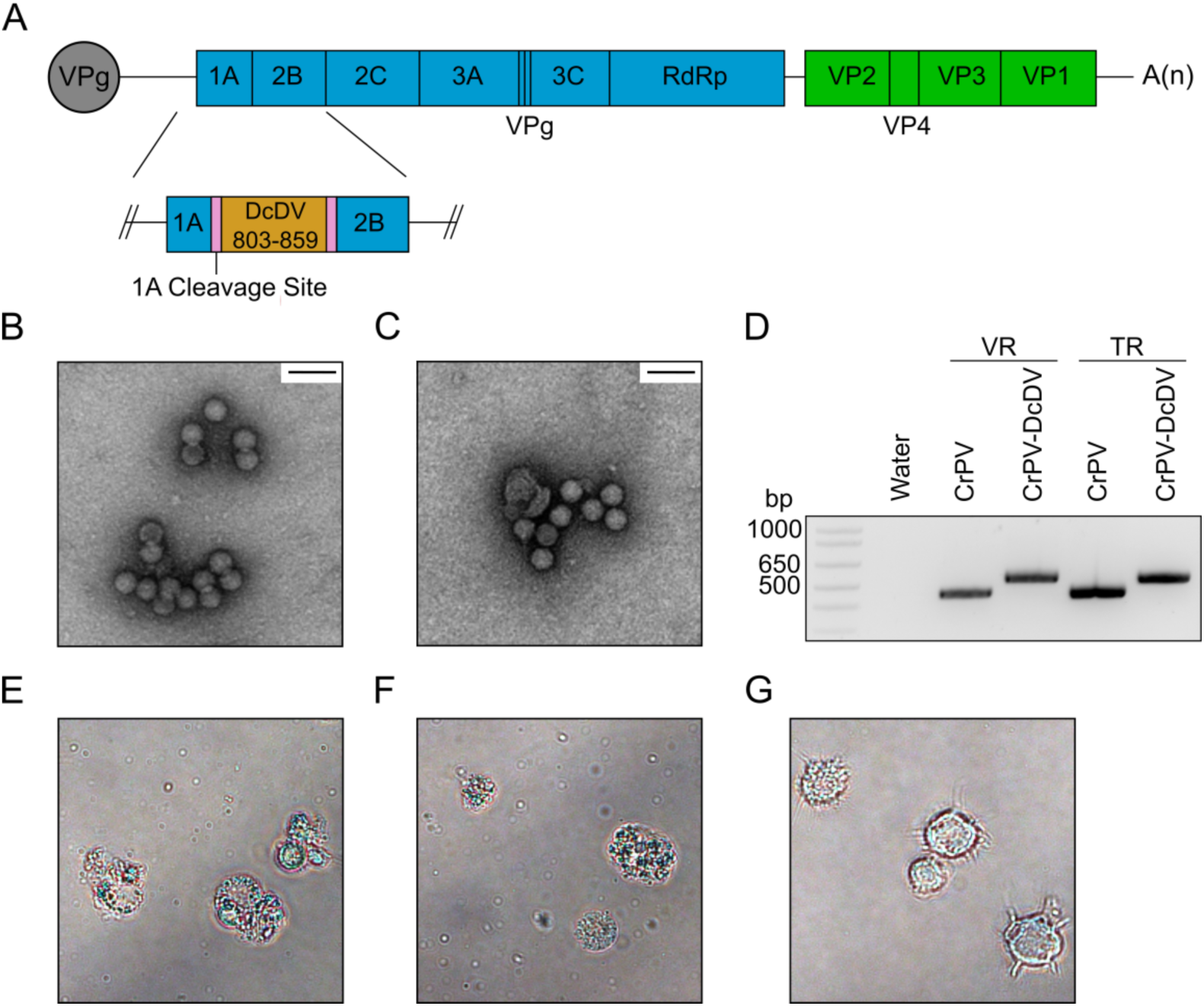
Construction of CrPV-DcDV, a recombinant CrPV mutant containing 57 nt of sequence from the DcDV genome. (A) Genome organization of CrPV DcDV. Blue rectangles represent the CrPV non-structural proteins. RdRp = RNA dependent RNA polymerase. VPg = viral protein genome-linked. Green boxes represent the CrPV structural proteins. The orange rectangle represents the recombinant DcDV sequence, which corresponds to nucleotides 803-859 from the DcDV genome. Pink rectangles represent the cleavage site at which the 1A protein is released from the polyprotein. (B & C) Electron micrographs of wild-type CrPV (B) or CrPV-DcDV (C) virions purified from S2 cells transfected with viral RNA. 50,000x magnification. Scale bar is 50 nm. (D) RT-PCR products produced using primers flanking the site into which recombinant DcDV sequence was inserted in CrPV-DcDV (primers 11 and 12). RNA extracted from purified virions (VR) or *in vitro* transcribed viral RNA (TR) was used as a template. (E-G) Bright-field microscopy images of S2 cells infected with wild-type CrPV virions (E), CrPV-DcDV virions (F), or mock infected (G). Images were acquired 72 hours post-infection.

We found that the virions produced following transfection of S2 cells with *in vitro* transcribed CrPV-DcDV RNA were indistinguishable from those produced following transfection with wild-type CrPV RNA (Fig. 5b & c) and RT-PCR of RNA purified from CrPV-DcDV virions indicated that the recombinant sequence was retained (Fig. 5d). Finally, we found that infection of S2 cells with either wild-type CrPV or CrPV-DcDV virions resulted in the cytopathic effects characteristic of CrPV infection (Fig. 5e & f).

To determine whether ENS-derived piRNAs mediate a response to CrPV-DcDV in *D. citri*, we inoculated CRF-CA *D. citri* with 1000 tissue culture infectious dose 50% (TCID_50_) units of wild-type CrPV or CrPV-DcDV per insect by intrathoracic injection. We observed a significant difference in viral RNA levels for wild-type CrPV and CrPV-DcDV three days post injection (Fig. 6a). By five days post injection, CrPV-DcDV RNA levels remained lower than those of wild-type CrPV, but the difference was not statistically significant (Fig. 6a). RT-PCR results indicated the recombinant DcDV sequence was retained throughout the course of infection with CrPV-DcDV (Fig. S9). To determine if CrPV-DcDV was targeted by piRNAs, we sequenced sRNAs from insects collected five days post infection. During infection with either virus, we observed an abundance of 21 nt sRNAs of both polarities and these 21 nt sRNAs mapped to the entire sequence of both viral genomes, indicating the presence of an siRNA-based response (Fig. 6b & c, Fig. S10 a & b). The recombinant DcDV sequence within the CrPV-DcDV genome was a target of this siRNA-based response, as we observed abundant 21 nt sRNAs derived from this portion of the recombinant virus genome in *D. citri* infected with CrPV-DcDV (Fig. 6d). When sRNAs from *D. citri* infected with wild-type CrPV were mapped to the same DcDV-derived sequence, a peak within the piRNA-size range was observed due to the production of piRNAs from ENS, but no 21 nt peak was seen (Fig. 6e)

**Fig. 6.**
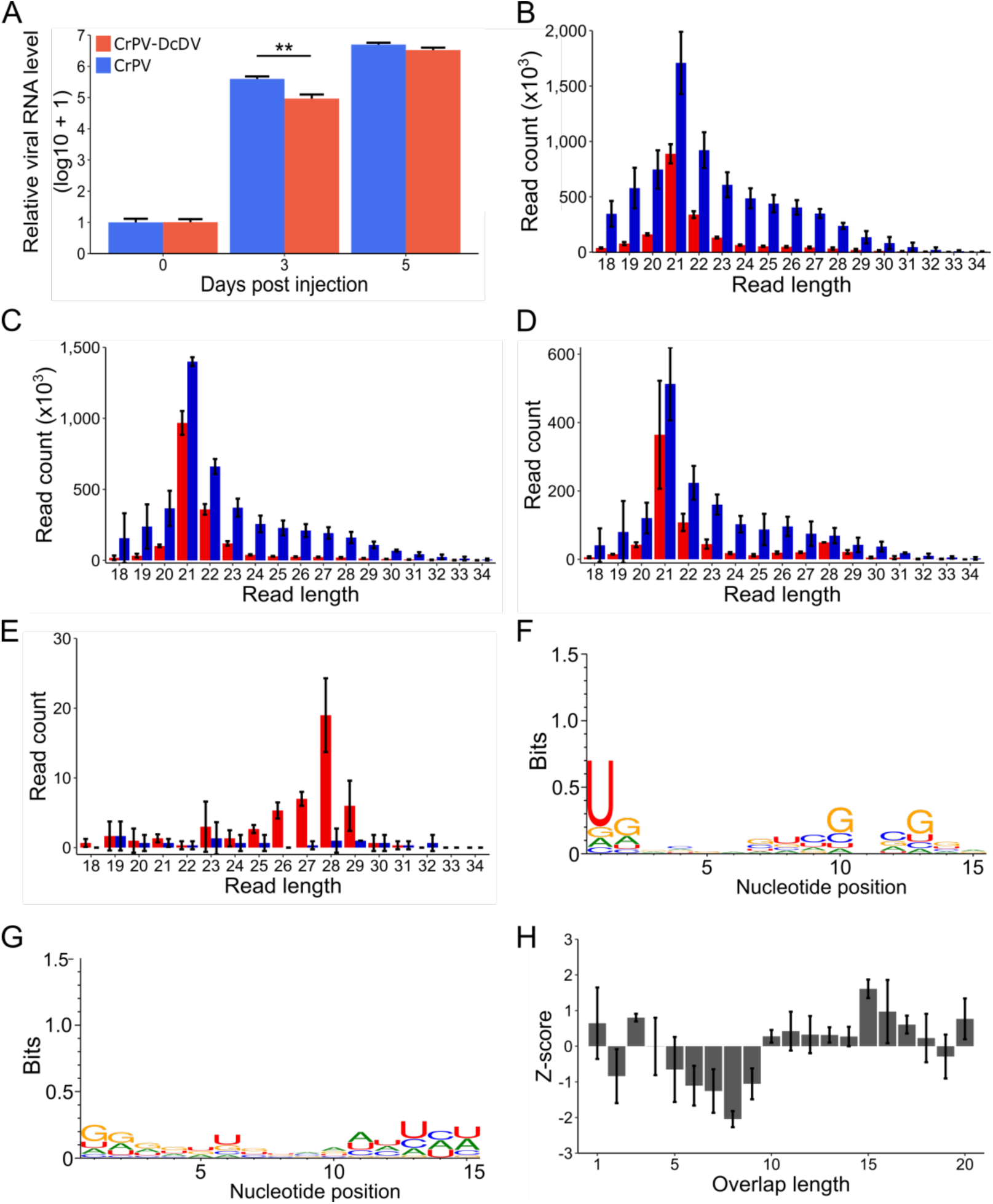
CrPV-DcDV is not targeted by piRNAs during infection initiated by intrathoracic injection of purified virions. (A) Relative viral RNA levels during infection with wild-type CrPV or CrPV-DcDV. Infection was initiated by intrathoracic injection of CRF-CA *D. citri* with 1000 TCID_50_ units of purified virions per insect. Viral RNA levels were assessed by RT-qPCR and normalized based on the expression of actin. The amount of viral RNA present on day 0 was set as 1 and log10 + 1 levels of viral RNA are shown relative to this value. Bars represent the average viral RNA level in five pools of three insects. Error bars indicate the standard error of the mean. ** = p < 0.01, two-tailed T-test. (B-F) analysis of sRNA sequencing data of sRNAs purified from CRF-CA *D. citri* infected with wild-type CrPV or CrPV-DcDV by intrathoracic injection as described for (A). sRNA was purified from pools of 25 *D. citri* collected 5 days post injection. (B) Length distribution of sRNAs from wild-type CrPV-infected *D. citri* mapped to the wild-type CrPV genome. Red = antisense, blue = sense. Read counts are an average of three independent libraries. Error bars indicate standard deviation. (C) Length distribution of sRNAs from CrPV-DcDV-infected *D. citri* mapped to the CrPV-DcDV genome. Red = antisense, blue = sense. Read counts are an average of three independent libraries. Error bars indicate standard deviation. (D) Length distribution of sRNAs from CrPV-DcDV-infected *D. citri* mapped to the recombinant DcDV sequence present within the CrPV-DcDV genome. Red = antisense, blue = sense. Read counts are an average of three independent libraries. Error bars indicate standard deviation. (E) Length distribution of sRNAs from wild-type CrPV-infected *D. citri* mapped to the recombinant DcDV sequence present within the CrPV-DcDV genome. Red = antisense, blue = sense. Read counts are an average of three independent libraries. Error bars indicate standard deviation. (F & G) Sequence logos for 27-32 nt sRNAs from CrPV-DcDV-infected *D. citri* mapped to the recombinant DcDV sequence present within the CrPV-DcDV genome. Sequence logos for antisense sRNAs (F) or sense sRNAs (G) are shown. Data represents three pooled libraries. (H) Probability of overlap between the 5’ ends of complementary 27-32 nt sRNAs mapping to opposite strands of the recombinant DcDV sequence present within the CrPV-DcDV genome during infection with CrPV-DcDV. Probabilities are shown for the indicated overlap distances and represent the average of three independent libraries. Error bars indicate standard deviation. The average Z-score and the standard deviation of the Z-score for an overlap length of 10 nt is shown

To evaluate whether CrPV-DcDV was targeted by ping-pong-dependent piRNAs, we analyzed 27-32 nt sRNAs separately depending on whether they mapped to the recombinant DcDV sequence within the CrPV-DcDV genome or to the rest of the CrPV genome. We detected ENS-derived antisense piRNAs mapping to the recombinant DcDV sequence in *D. citri* infected with either wild-type CrPV or CrPV-DcDV (Fig. 6d-f), however, there was no evidence for the production of ping-pong-dependent secondary piRNAs from this region during infection with CrPV-DcDV (ping-pong Z-score = 0.28±0.18)(Fig. 6g & h). Ping-pong signatures were also not observed for the non-recombinant portion of the CrPV-DcDV genome or for wild-type CrPV (Fig. S10 c & d). These results indicate that CrPV-DcDV is not targeted by piRNAs during infection initiated by intrathoracic injection in *D. citri* despite the presence of ENS-derived piRNAs identical to the recombinant DcDV sequence.

Because the expression of ENS-derived piRNAs is highest in the gut (Fig. 3d) we wanted to examine whether a piRNA-based response to CrPV-DcDV could be detected during infection initiated by oral acquisition. Thus, we orally inoculated *D. citri* insects from CRF-CA with wild-type CrPV or CrPV-DcDV by allowing the insects to feed for 96 hours on an artificial diet solution containing 10^9^ TCID_50_ units/mL of wild-type CrPV or CrPV-DcDV. Following the 96 hour feeding period, the insects were transferred to *Citrus macrophylla* plants and viral RNA levels were evaluated every three days by RT-qPCR. Viral RNA levels in the insects were nearly identical for wild-type CrPV and CrPV-DcDV immediately following their removal from the virus-containing artificial diet and average viral RNA levels increased in the days following transfer of the insects to plants, although there was a substantial amount of variation between biological replicates (Fig. 7a). As with infections initiated by intrathracic injection, RT-PCR results indicated the recombinant DcDV sequence was retained throughout the course of oral infection with CrPV-DcDV (Fig. S9b). These results suggest that while wild-type CrPV and CrPV-DcDV are both infectious in *D. citri* following oral acquisition, individual insects displayed a variety of infection outcomes. Similar results have previously been reported for orally acquired infections with other RNA viruses in insects (45). We sequenced sRNAs from insects collected nine days after transfer of the insects from the virus-containing artificial diet to plants. We analyzed this data as described above for infections initiated by injection and found that while both wild-type CrPV and CrPV-DcDV were targeted by siRNAs, neither virus was targeted by ping-pong-dependent vpiRNAs during oral infection (ping-pong Z-score for 27-32 nt sRNAs mapping to the recombinant DcDV sequence during CrPV-DcDV infection = 1.04 ± 0.67)(Fig. 7b-h). Together, our results indicate that despite the presence of endogenous primary piRNAs complementary to a portion of the recombinant viral genome, CrPV-DcDV is not a target of the piRNA pathway in *D. citri* following infection initiated either orally or by intrathoracic injection.

**Fig. 7.**
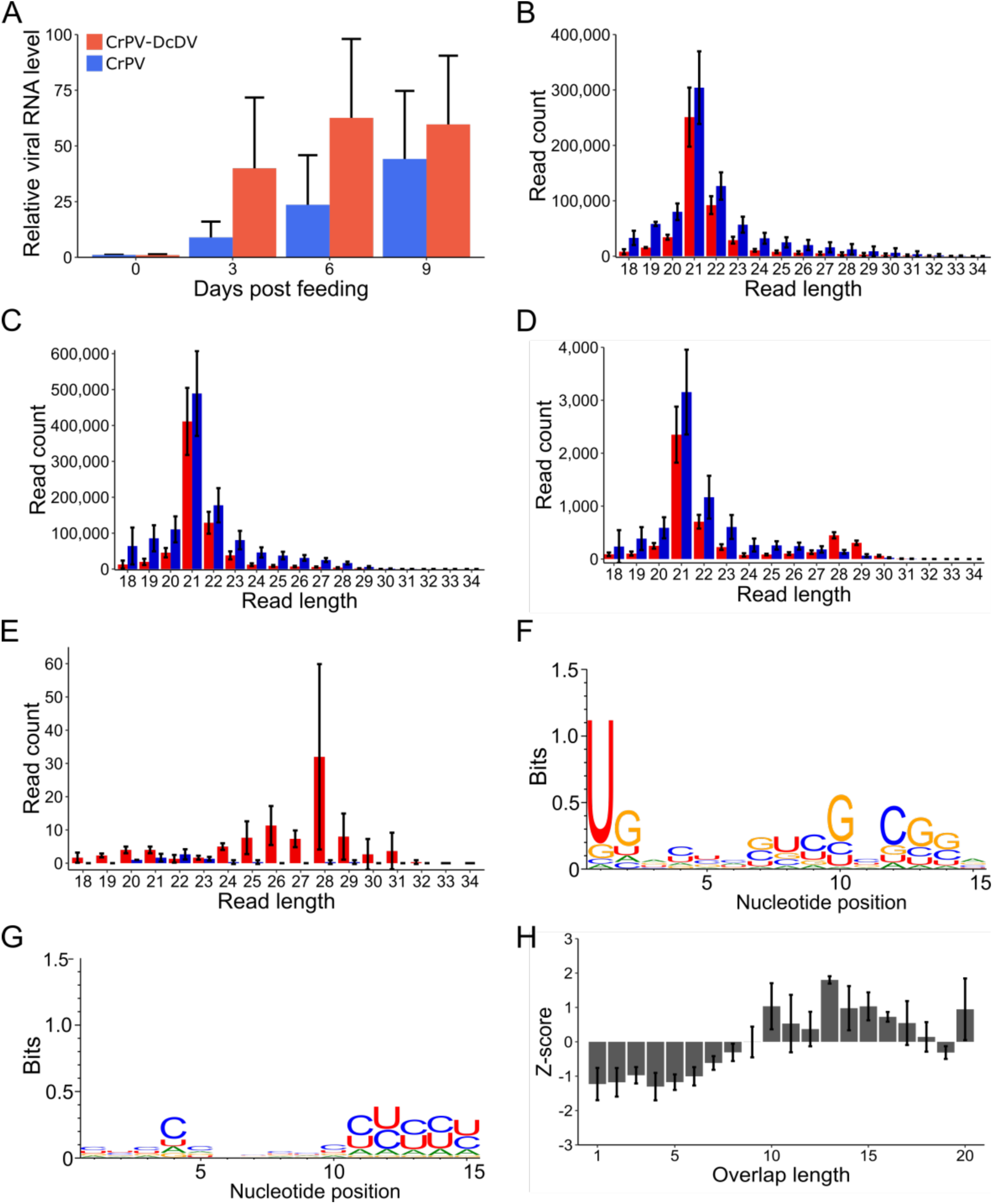
CrPV-DcDV is not targeted by piRNAs during infection initiated by oral acquisition of purified virions. (A) Relative viral RNA levels during infection with wild-type CrPV or CrPV-DcDV. Infection was initiated in CRF-CA *D. citri* by oral acquisition. Insects were allowed to feed for 96 hours on an artificial diet solution containing 10^9^ TCID_50_ units/mL of wild-type CrPV or CrPV-DcDV. Following the feeding period insects were moved to *C. macrophylla* plants (day 0 post feeding). Viral RNA levels were assessed by RT-qPCR and normalized based on the expression of actin. The amount of viral RNA present on day 0 was set as 1 and viral RNA levels are shown relative to this value. Bars represent the average viral RNA level in seven pools of three insects. Error bars indicate the standard error of the mean. (B-F) analysis of sRNA sequencing data of sRNAs purified from CRF-CA *D. citri* infected with wild-type CrPV or CrPV-DcDV by oral acquisition as described for (A). sRNA was purified from pools of 25 *D. citri* collected 9 days post feeding. (B) Length distribution of sRNAs from wild-type CrPV-infected *D. citri* mapped to the wild-type CrPV genome. Red = antisense, blue = sense. Read counts are an average of three independent libraries. Error bars indicate standard deviation. (C) Length distribution of sRNAs from CrPV-DcDV-infected *D. citri* mapped to the CrPV-DcDV genome. Red = antisense, blue = sense. Read counts are an average of three independent libraries. Error bars indicate standard deviation. (D) Length distribution of sRNAs from CrPV-DcDV-infected *D. citri* mapped to the recombinant DcDV sequence present within the CrPV-DcDV genome. Red = antisense, blue = sense. Read counts are an average of three independent libraries. Error bars indicate standard deviation. (E) Length distribution of sRNAs from wild-type CrPV-infected *D. citri* mapped to the recombinant DcDV sequence present within the CrPV-DcDV genome. Red = antisense, blue = sense. Read counts are an average of three independent libraries. Error bars indicate standard deviation. (F & G) Sequence logos for 27-32 nt sRNAs from CrPV-DcDV-infected *D. citri* mapped to the recombinant DcDV sequence present within the CrPV-DcDV genome. Sequence logos for antisense sRNAs (F) or sense sRNAs (G) are shown. Data represents three pooled libraries. (H) Probability of overlap between the 5’ ends of complementary 27-32 nt sRNAs mapping to opposite strands of the recombinant DcDV sequence present within the CrPV-DcDV genome during infection with CrPV-DcDV. Probabilities are shown for the indicated overlap distances and represent the average of three independent libraries. Error bars indicate standard deviation. The average Z-score and the standard deviation of the Z-score for an overlap length of 10 nt is shown.

## Discussion

Primary vpiRNAs were first reported for American nodavirus, Drosophila C virus, Drosophila tetravirus, Nora virus, Drosophila X virus, and Drosophila birnavirus in a *D. melanogaster* ovarian somatic sheet (OSS) cell line (46). However, these conclusions were based only the size of the sRNAs and nucleotide biases for all virus-specific sRNAs pooled into a single analysis. A subsequent study using whole flies found no evidence for the production of primary or secondary vpiRNAs during infection with Drosophila C virus, Drosophila X virus, Sindbis virus, Nora virus, Drosophila A virus, Drosophila melanogaster sigma virus, invertebrate iridescent virus 6, or Flock house virus (14). To date, ping-pong dependent vpiRNAs have only been found in a small number of mosquito species and cells lines that express an expanded group of Piwi-family Argonaute proteins.

Aedes albopictus densovirus 1 (AalDV-1) was recently shown to be a target of both siRNAs and ping-pong-dependent vpiRNAs during persistent infection in *Ae. aegypti*-derived Aag2 cells, representing the first report of vpiRNAs targeting a DNA virus and the first report of siRNAs targeting a ssDNA virus in insects (47). Densoviruses replicate exclusively in the nucleus (48) and interestingly, analysis of nuclear and cytoplasmic sRNA fractions of AalDV-1-infected Aag2 cells revealed that while AalDV-1-derived primary piRNAs are found in both the nucleus and cytoplasm, ping-pong-dependent piRNAs targeting AalDV-1 are found only in the cytoplasm (47). Ago3 and Piwi5 are the sole Piwi family proteins required for the production of vpiRNAs during infection of Aag2 cells with Sindbis virus (9). As these proteins are expressed only in the cytoplasm, the observation that AalDV-1-derived primary piRNAs are present in the nucleus during infection of Aag2 cells suggests that a different pathway may generate primary vpiRNAs from viral RNA in the nucleus (47). Such a pathway would be distinct from the cytoplasmic Ago3/Piwi5 pathway that is responsible for production of vpiRNAs from Sindbis virus and might rely on a piRNA biogenesis factor expressed in the nucleus, such as zucchini endonuclease (47).

How viruses are marked as substrates for piRNA biogenesis remains unclear. In Aag2 cells, viral and TE RNAs are processed into piRNAs by distinct sets of Piwi-family Argonaute proteins, but it is unknown how viral RNA is recognized by these proteins (9). Moreover, as discussed above, the presence of AalDV-1-derived primary piRNAs in the nucleus of Aag2 cells suggests that a different pathway for the production of primary vpiRNAs may operate in the nucleus (47). Several previous reports have speculated that primary piRNAs derived from EVEs may facilitate the targeting of cognate viruses in a manner analogous to the targeting of TEs by the canonical ping-pong cycle (15–17). Theoretically, such a targeting mechanism could facilitate the production of vpiRNAs in species that do not express the expanded group of Piwi-family Argonaute proteins expressed by some mosquito species and mosquito-derived cell lines.

The *D. citri* genome contains a DcDV-derived EVE (denoted ENS) with 86% nucleotide identity to the corresponding portion of the DcDV genome and here we found that ENS serves as a source for DcDV-specific primary piRNAs in geographically distinct *D. citri* populations. However, we found that ENS is not present in all *D. citri* populations and that *D. citri* insects lacking ENS do not produce endogenous DcDV-specific piRNAs. While more comprehensive analyses are needed, comparing our results with the known geographic distribution of *D. citri* lineages suggests that ENS may be present in lineage B *D. citri* and absent in lineage A *D. citri.* We previously established a colony of *D. citri* insects harboring DcDV as a persistent infection (CRF-TW)(34) and here we found that insects from this colony do not possess ENS. Sequencing of sRNAs from *D. citri* insects from CRF-TW revealed the production of DcDV-derived ping-pong-dependent vpiRNAs. Similar results were obtained by sequencing sRNAs from DcDV-infected CRF-Uru *D. citri*, another *D. citri* population which does produce any endogenous DcDV-specific piRNAs. These were surprising results, as the *D. citri* repertoire of Piwi-family Argonaute proteins is limited to homologs of those expressed in *D. melanogaster,* which have so far not been associated with the production of vpiRNAs (29, 49). The production of vpiRNAs in *D. citri* seems to display some virus specificity as vpiRNAs were not seen during infection with Diaphorina citri reovirus, Diaphorina citri picrona-like virus, Diaphorina citri bunyavirus, Diaphorina citri flavi-like virus, or CrPV. Notably, these viruses are all RNA viruses that replicate in the cytoplasm, but DcDV is a DNA virus that replicates in the nucleus. While the production of ping-pong-dependent DcDV-derived vpiRNAs is noteworthy, we cannot determine whether this process plays an antiviral role based on the present data.

Densoviruses are known to replicate exclusively in the nucleus (48). Given that vpiRNAs are not produced during infection of *D. citri* with positive strand RNA viruses, a negative strand RNA virus, or a dsRNA virus, future analysis should determine whether DcDV RNA is processed into primary vpiRNAs in the nucleus as was suggested for AalDV-1 (47). Interestingly, abundant virus-derived sRNAs within the piRNA size range are also observed during infection of *Myzus persicae* with Myzus persicae densovirus and during infection of *Culex pipiens molestus* with Mosquito densovirus, however, these sRNAs were not analyzed for the presence of piRNA signatures (50, 51).

In addition to ping-pong dependent vpiRNAs, our results show that DcDV is also targeted by 21 nt siRNAs. RNA interference mediated by siRNAs is the primary antiviral defense mechanism against RNA viruses in insects and requires cleavage of dsRNA substrates by the RNase III enzyme dicer (52). RNA interference mediated by siRNAs is also known to target dsDNA viruses and a ssDNA virus was recently shown to be targeted by siRNAs in Aag2 cells (47,53– 56). Previous reports have suggested that overlapping viral transcripts represent the dsRNA substrate that becomes processed into siRNAs during infection with DNA viruses (47,53–55). Indeed, we previously found that transcriptional readthrough during transcription of the ambisense DcDV genome leads to the production of nearly genome length complementary transcripts (34). These transcripts may anneal to form the dsRNA required for production of siRNAs and may constitute the sense and antisense precursors required for the ping-pong cycle.

Although our results indicate that EVE-derived piRNAs are not required for the production of ping-pong-dependent DcDV-derived vpiRNAs, we cannot exclude the possibility that EVE-derived piRNAs could serve as an additional pool of piRNAs that could also contribute to priming of the ping-pong cycle. Thus, to investigate whether EVE-derived piRNAs can prime ping-pong amplification during infection with a virus sharing complementary sequence in *D. citri*, we constructed CrPV-DcDV, a recombinant CrPV harboring 57 nt derived from the DcDV genome and corresponding to ENS. Despite the presence of endogenous primary piRNAs derived from ENS perfectly mapping to the recombinant region of CrPV-DcDV, we did not observe induction of the ping-pong cycle during infection with this virus initiated either by oral acquisition or intrathoracic injection of virions. The lack of ping-pong cycle activation during infection with CrPV-DcDV is unlikely to be due to an absence of ping-pong activity in the tissues infected by CrPV-DcDV, as we detected abundant TE-derived ping-pong-dependent piRNAs in dissected *D. citri* guts, heads, hemolymph, ovaries, and testis. These results indicate that the presence of EVE-derived primary piRNAs sharing perfect nucleotide identity with an exogenous virus is not in itself sufficient to mark that virus as a target for ping-pong amplification in *D. citri*. We note that our approach has some limitations. Host-responses to virus infection are shaped by coevolution and there is a possibility that CrPV-DcDV was not targeted by ping-pong-derived piRNAs due to differences in the infection cycles between DcDV (a naturally infecting DNA virus) and CrPV (a non-naturally infecting RNA virus). For example, these may include potential differences in the spatial or physical accessibility of viral RNA to piRNA biogenesis factors. Indeed, our results suggest that RNA viruses are not targeted by piRNAs in *D. citri*. Thus, it is possible that even if EVE-derived piRNAs could prime ping-pong amplification during infection with a cognate DNA virus, cognate RNA viruses may not be susceptible to such targeting by the piRNA pathway. Future research aiming to uncover the mechanisms of vpiRNA production during infection with DNA viruses such as DcDV and AalDV-1 will help to resolve these questions.

We previously reported that while DcDV persistently infects CRF-TW *D. citri,* CRF-CA *D. citri* are not susceptible to DcDV infection by oral acquisition or intrathoracic injection (34). We report here that CRF-Uru *D. citri* can be persistently infected with DcDV by either of these routes of infection. As discussed above, CRF-TW and CRF-Uru *D. citri* are likely to be of lineage A, while CRF-CA *D. citri* are likely to be of lineage B. At present, no genotypic or phenotypic differences between the two lineages have been described except for nucleotide variation within the mitochondrial cytochrome oxidase subunit I gene (30–32). Genetic background is known to contribute to susceptibility to virus infection within different populations of the same species (57). In the case of Densoviruses, a single arginine residue at position 188 within a mucin-like glycoprotein expressed in the midgut epithelium of *Bombyx mori* strains susceptible to infection with Bombyx mori densovirus (BmDV) was substituted by other amino acids in resistant strains (58). Further analysis showed that this mucin-like glycoprotein was the cellular receptor for BmDV in the midgut epithelium and that the arginine residue at position 188 was required for BmDV binding to this protein (58). EVEs have been suggested to play a role in susceptibility to virus infection through a variety of potential mechanisms (16,59–61). Our results suggest that ENS is present in lineage B *D. citri,* but absent in lineage A *D. citri* and that the presence of ENS correlates with resistance to DcDV infection in the populations tested. However, other genetic differences between the two lineages are likely to exist and we cannot assess whether ENS may be involved with resistance to DcDV infection based on the present data. Nevertheless, our data highlight both a genetic and phenotypic difference between the two *D. citri* lineages. Other such differences between the lineages and how they may contribute to susceptibility to DcDV infections should be the subject of future investigations.

## Materials and methods

### Maintenance of *D. citri* insects

*D. citri* insects were reared on *C. macrophylla* plants in mesh cages (BugDorm, Taichung, Taiwan) at 25 ± 2 °C under a 14:10 hour (light:dark) photoperiod and 60-70% relative humidity at the University of California Davis CRF (62). *D. citri* insects from Uruguay and Taiwan were imported under USDA APHIS-PPQ permit P526P-17-02906 by shipping adults and nymphs on *C. macrophylla* or *Murraya paniculata* cuttings, respectively.

### sRNA sequencing and analysis

For total insect sRNA sequencing of CRF-CA, CRF-TW, and CRF-Uru *D. citri*, total RNA was extracted from groups of 50 adult *D. citri* insects using TRIzol reagent (Invitrogen, Carlsbad, California) according to the manufacturer’s instructions, but omitting the 75% ethanol wash. For CRF-CA *D. citri* infected with wild-type CrPV or CrPV-DcDV by oral acquisition or injection and for the progeny CRF-Uru *D. citri* infected with DcDV by injection, sRNAs were extracted from groups of 25 adult *D. citri* insects using TRIzol reagent according to the manufacturer’s instructions, but omitting the 75% ethanol wash. For organ specific sRNA sequencing multiple dissected organs were combined to be processed as one sample: 200 for gut, head, or hemolymph and 100 for ovary or testis. RNA extraction was done from pooled organ samples with the Direct-zol RNA Miniprep kit (Zymo Research, Irvine, California) following the manufacturer’s instructions. Three pooled samples of each organ were used. For all sRNA sequencing, total RNA was sent to Beijing Genomics Institute for library prep with the TruSeq small RNA sample preparation kit (Illumina, San Diego, California) and sequencing by 50 bp single end sequencing with the BGISEQ-500 platform. For all sRNA reads, adaptor sequences were removed with Trim Galore version 0.4.4 (63) and the fastxtoolkit was used to remove reads containing bases with quality score <20 (64). The remaining reads were mapped to viral genomes, insect genomes, or TEs using BowTie version 1.2.1.1 (65) using the default settings with the following exceptions: - n 1 -l 20. When only perfectly mapped reads were considered (specifically indicated in the text), the –v 0 option was used instead of the –n 1 –l 20 options. Sequence logos were produced using WebLogo 3 (66). Ping-pong Z-scores were calculated with signature.py (67). All sRNA sequence data generated for this study was deposited to the NCBI SRA database (BioProject accession no. PRJNA629895).

### Analysis of EVEs

DcDV-derived EVEs were identified in *D. citri* genomic scaffold 2850 by BLASTx using as queries the deduced amino acid sequences of DcDV structural and non-structural proteins (GenBank accession no.’s YP_009256210, YP_009256211, YP_009256212, and YP_009256213). DcDV-derived EVEs were then inspected by aligning the DcDV genome (GenBank accession no. NC_030296) to the nucleotide sequence of *D. citri* genomic scaffold 2850 using ClustalW (68). Deduced amino acid and nucleotide identities shared between ENS and DcDV and between EITR and DcDV were calculated as p-distance with complete deletion of gaps from ClustalW alignments using MEGA 7 (69).

### Identification of TEs

We identified TEs present within the *D. citri* genome using RepeatMasker version 4.0.6 (70) with the Metazoa library. In addition, to identify TEs lacking homology to previously annotated TEs, we used RepeatModeler version 1.0.8 (71) to produce a *de novo* hidden Markov model for TEs within the *D. citri* genome which was subsequently used as input for a second analysis using RepeatMasker. All TE identification was performed using TEAnnotator.py as previously described to produce a single strand-specific .fasta file containing the sequences of all TEs >100 nt identified in the *D. citri* genome (35).

### Nucleic acid extraction, reverse transcription, PCR, exonuclease digestion, and Sanger sequencing

Unless otherwise specified, DNA was extracted from groups of 25 homogenized *D. citri* insects by phenol/chloroform extraction followed by ethanol precipitation. For PCR using exonuclease treated DNA, 1 μg CRF-CA *D. citri* genomic DNA, an intact plasmid containing the ENS amplicon, or the same plasmid digested overnight with HindIII-HF and SbfI-HF (New England Biolabs, Ipswich, Massachusetts) were digested overnight with 10 units of Plasmid-Safe ATP-Dependent DNAse (Lucigen, Middleton, Wisconsin) according to the manufacturer’s instructions. For analysis of ENS transcription, RNA was extracted from groups of 25 homogenized CRF-CA *D. citri* using TRIzol reagent according to the manufacturer’s instructions. RNA was treated twice with the Qiagen RNAse-free DNAse set (Qiagen, Hilden, Germany) to remove DNA. cDNA was prepared from 50 ng RNA with SuperScript IV reverse transcriptase (Invitrogen, Carlsbad, California) using specific primers (see the legend for Fig. 1) according to the manufacturer’s instructions. To assess retention of the insert in CrPV-DcDV, RNA was extracted from CrPV or CrPV-DcDV virions using TRIzol LS (Invitrogen, Carlsbad, California) according to the manufacturer’s instructions. cDNA was prepared from 50 ng virion RNA using the Applied Biosystems high capacity reverse transcription kit (Applied Biosystems, Foster City, California) using random primers according to the manufacturer’s instructions and cDNA was diluted 1:10 prior to PCR. Primer sequences are given in supplementary Table 1 and their use is described in the appropriate figure legends. All PCR reactions were performed with CloneAmp HiFi PCR premix (Takara Bio, Mountain View, California) using 25 ng of DNA, 1 µl of cDNA (for determination of ENS transcription), or 3 µl diluted cDNA (for determination of CrPV-DcDV insertion retention). Detailed PCR protocols are available upon request. PCR products were analyzed by 0.8% or 2% agarose gel electrophoresis and visualized under UV after staining with SYBR safe DNA gel stain (Invitrogen, Carlsbad, California). PCR products were purified with the Zymoclean Gel DNA Recovery Kit (Zymo Research, Irvine, California) and sequenced by Quintarabio using the Sanger method.

### Southern blotting

5 µg undigested CRF-CA *D citri* DNA or 5 µg CRF-CA *D. citri* DNA digested overnight with PstI-HF and HindIII-HF (New England Biolabs, Ibswich, Massachusetts) was electrophoresed in a 0.8% agarose/0.5x TAE gel and the gel was prepared for transfer by incubation in 0.25 M HCl for 30 minutes, then 0.5 M NaCl, 0.5 M NaOH for 30 minutes, and then 1.5 M NaCl, 0.5 M Tris-HCl pH 7 for 30 minutes. DNA was then transferred to an Amersham Hybond Nx membrane (GE Life Sciences, Boston, Massachusetts) for 24 hours in 20X SSC using the capillary method. DNA was cross-linked to the membrane by exposure to 120,000 µjoles of UV light using a UV Stratalinker 2400 (Stratagene, San Diego, California).

A PCR product corresponding to ENS was prepared by PCR with primers 27 and 3 using CRF-CA *D. citri* DNA as a template. This PCR product was then used a template to prepare a radioactive RNA probe by *in vitro* transcription using the MAXIscript T7 Transcription Kit (Invitrogen, Carlsbad, California) with 32P-UTP according to the manufacturer’s instructions.

The membrane with cross-linked DNA was prehybridized by incubation in Ambion ULTRAhyb Ultrasensitive hybridization buffer (Invitrogen, Carlsbad, California) for 2 hours at 42 °C in a rotary incubation oven. Following prehybridization, 2 x 10^6^ CPM/mL radioactive probe was added and hybridization proceeded for 16 hours at 42 °C in a rotary incubation oven. The membrane was then washed at 42 °C for 15 minutes each with 2X SSC/0.1% SDS, 0.2X SSC/0.1% SDS, and 0.1X SSC/0.1% SDS.

### DcDV infections and DcDV qPCR

DcDV virions were partially purified from 0.5 g of CRF-TW *D. citri* as previously described (34). The number of DcDV genome copies in the virion preparation was determined by qPCR with primers 29 and 30 (see table S1) as described previously (34). For infections by intrathoracic injection, the virion preparations were diluted to 8.2 x 10^5^ genome copies/µl in 100 mM Tris-HCl, pH 7.5. Adult CRF-Uru *D. citri* were anesthetized on ice for approximately 15 minutes and intrathoracicly injected with approximately 200 nl of the diluted virion preparation by manual injection using a syringe fitted to a 34 gauge stainless steel needle (Hamilton, Reno, Nevada). Following injection, insects were maintained on *C. macrophylla* plants. For infections by oral acquisition, the virion preparations were diluted to 8.2 x 10^5^ genome copies/µl in 15% sucrose, 0.1% green food coloring, 0.4% yellow food coloring prepared in 100 mM Tris-HCl, pH 7.5. Groups of 30 adult CRF-Uru *D. citri* were fed on the virus-containing sucrose solution for 96 hours by membrane feeding as described previously (34). Following the feeding period, insects were maintained on *C. macrophylla* plants. Day 0 is designated as the day insects were transferred from the virus containing artificial diet to plants. For both types of infection, adult insects (>1 day old) were used and neither age nor gender of the insects was determined prior to infection.

For both types of infection, five pools of three insects were collected every two days and DNA was purified from each pool by phenol/chloroform extraction followed by ethanol precipitation as described previously (34). All insects had been collected or were removed from the plants by 17 days post infection. During the maintenance period on plants, the DcDV-injected and -fed insects laid eggs on the plants. Following removal of the DcDV-injected or –fed insects, the plants were maintained to allow development of the progeny. The progeny were removed on the day of emergency to the adult stage and DNA was extracted from 10 individual progeny insects from each experiment as described above. A pool of 25 progeny of the DcDV-injected *D. citri* was collected at the same time for sRNA purification. The concentration of DcDV genome copies in the DNA pools or in DNA extracted from individual progeny insects was assessed by qPCR with primers 29 and 30 (see Fig. S1) using 25 ng DNA as described previously (34). Technical triplicates were used for all qPCR reactions.

### Cloning of CrPV-DcDV

Complementary lyophilized oligonucleotides containing the DcDV sequence to be inserted flanked upstream by 15 nt corresponding to the 3’ end of the CrPV 1A nucleotide sequence and downstream by 15 nt corresponding to the 5’ end of the CrPV 2B nucleotide sequence were suspended to a concentration of 100 µM in 100 mM potassium acetate, 300 µM HEPES, pH 7.5 (oligo sequences: 5’-TCTAATCCTGGTCCTGCAGACCGTTCACCTTCTCCAGGACCTTCTACTGCATATCGCTATTGTAGCGAGGAAGTGCAATCGCGCCCC and 5’ – GGGGCGCGATTGCACTTCCTCGCTACAATAGCGATATGCAGTAGAAGGTCCTGGAGAAGGTGAACGGTCTGCAGGACCAGGATTAGA). To anneal these oligonucleotides, the oligonucleotides were mixed in equimolar ratios, incubated at 94 °C for 3 minutes, and allowed to cool slowly to room temperature to form the duplex oligo. pCrPV-3 (a gift from Shou-Wei Ding) (72) was linerized by inverse PCR with primers 13 and 14 and the duplex oligo was ligated to the linearized plasmid in a 2:1 insert:vector molar ratio using NEBuilder Hifi Assembly Master Mix (New England Biolabs, Ipswich, Massachusetts) according to the manufacturer’s instructions. The resulting plasmid (designated pCrPV-1A-DcDV) was transformed into chemically competent *E. coli* DH5α and transformed cells were grown on luria bertani (LB) agar plates containing 100 µg/mL ampicillin at 28 °C for 16 hours. Insert presence was verified by colony PCR with primers 15 and 17 and by Sanger sequencing of colony PCR products. Colonies containing the desired plasmid were used to inoculate 5 mL LB broth containing 100 µg/mL ampicillin and the cultures were incubated at 28 °C for 16 hours with 220 rpm shaking. Plasmids were purified from overnight cultures with the QIAprep Spin Miniprep Kit (Qiagen, Hilden, Germany) according to the manufacturer’s instructions.

To duplicate the 1A cleavage site on the 3’ end of the recombinant DcDV sequence, pCrPV-1A-DcDV was linearized by inverse PCR with primers 17 and 18 and the linearized plasmid was circularized by blunt end ligation with T4 DNA ligase (New England Biolabs, Ipswich, Massachusetts) according to the manufacturer’s instructions to create pCrPV-1A-DcDV-1A. The ligation product was transformed into chemically competent *E. coli* DH5α and plasmids were purified as described.

### Transfection and infection of S2 Cells and virion purification

pCrPV-3 and pCrPV-1A-DcDV-1A were linearized by digestion with BamHI-HF (New England Biolabs, Ipswich, Massachusetts). Wild-type and CrPV-DcDV RNA was prepared by *in vitro* transcription of 1 µg linearized plasmid using the mMESSAGE mMACHINE T7 transcription kit (Invitrogen, Carlsbad, California) according to the manufacturer’s instructions. 24 hours prior to transfection, 5 x 10^6^ S2 cells were seeded in 2 mL Schneider’s Drosophila media (Thermo Fisher, Waltham, Massachusetts) supplemented with 10% heat-inactivated fetal bovine serum in 9.6 cm^2^ wells of a tissue culture plate and incubated at 26 °C. The following day, cells were transfected with wild-type CrPV or CrPV-DcDV RNA using the TransMessenger transfection reagent (Qiagen, Hilden, Germany) according to the manufacturer’s instructions and the transfected cells were incubated at 26 °C. After 72 hours, 1 mL of transfected cells was transferred to a 50 mL tissue culture flask containing 3 x 10^6^ S2 cells/mL that had been passed 24 hours prior in 6 mL Schneider’s Drosophila media supplemented with 10% heat-inactivated fetal bovine serum. After incubation at 26 °C for 96 hours, virions were purified from the infected S2 cells by centrifugation at 8,000 RPM for 10 minutes at 4 °C using a Beckman GSA rotor (Beckman Coulter, Brea, California). The supernatant was then collected and centrifuged at 8,000 RPM for 30 minutes at 4 °C using a Beckman GSA rotor. The supernatant was collected and centrifuged through a 15% sucrose cushion (prepared in 10 mM Tris-HCl, pH 7.5) at 45,000 RPM for 45 minutes at 11 °C in a Beckman 70.1 Ti rotor (Beckman Coulter, Brea, California). The pellet was resuspended in 10 mM Tris-HCl, pH 7.5 and then filtered through a 0.22 µm filter.

The titer of purified CrPV-DcDV or CrPV virions was measured by end-point dilution. Briefly, 4 x 10^5^ S2 cells were seeded in 500 µL Schneider’s Drosophila media supplemented with 10% heat-inactivated fetal bovine serum in each well of 24 well plates. Cells were incubated for 24 hours at 26°C. After 24 hours cells were infected with tenfold serial dilutions of purified virions in 100 µL volumes. Dilutions over the range of 10^-5^ to 10^-10^ were used and six individual wells were used for each dilution. After 72 hours, cells were examined for the presence cytopathic effects and the titer of the undiluted virus stock was calculated from these results using the Reed and Muench method.

For infection of S2 cells using purified virions, 5 x 10^6^ S2 cells were seeded in 2 mL Schneider’s Drosophila media supplemented with 10% heat-inactivated fetal bovine serum in 9.6 cm^2^ wells of a tissue culture plate and incubated at 26 °C. After 24 hours, 100 TCID_50_ units of wild-type CrPV or CrPV-DcDV virions suspended in 100 µL Schneider’s Drosophila media were added to the wells.

### Microscopy

Wild-type CrPV or CrPV-DcDV virions (see above) were further purified by centrifugation through a 1.2 g/cm^3^ to 1.6 g/cm^3^ cesium chloride density gradient at 50,000 RPM for 4 hours at 11 °C in a Beckman SW 65 Ti rotor (Beckman Coulter, Brea, California). Visible “virus bands” in the gradients were collected using a Hamilton syringe and diluted 1:10 in 10 mM Tris-HCl, pH 7.5. Diluted virions were then centrifuged at 75,000 RPM for 30 minutes at 11 °C in a Beckman TLA 120 rotor (Beckman Coulter, Brea, California). The pellet was resuspended in 10 mM Tris-HCl, pH 7.5 and negatively stained with uranyl formate. Negatively stained virion samples were observed by transmission electron microscopy using a JEOL 1230 electron microscope (JEOL, Tokyo, Japan) operating at 100 kV. Mock-infected S2 cells and S2 cells infected with wild-type CrPV or CrPV-DcDV were observed by bright-field microscopy using a Leica DM5000B microscope (Leica, Wetzlar, Germany).

### CrPV infections in *D. citri*

Adult CRF-CA *D. citri* were infected with purified CrPV or CrPV-DcDV virions by intrathoracic injection or membrane feeding exactly as described above for DcDV infections. For injections, purified virions were diluted to 5000 TCID_50_ units/µl and insects were injected with approximately 200 nl each. Seven pools of three insects each were collected on days 0, 3, 6, and 9 for RNA extraction. Three additional pools of 25 insects each were collected on day 9 for sRNA extraction. For membrane feeding, purified virions were diluted to 10^9^ TCID_50_ units/µl. Five pools of three insects each were collected on days 0, 3, and 5 for RNA extraction. Three additional pools of 25 insects each were collected on day 5 for sRNA extraction. To determine viral RNA levels, RNA was extracted from the pools of three insects using TRIzol reagent according to the manufacturer’s instructions. RNA pellets were resuspended in 25 µl water and cDNA was prepared from 1 µl RNA (approximately 200-400 ng RNA) with the high capacity cDNA reverse transcription kit using random primers according to the manufacturer’s instructions and cDNA was diluted 1:10. Wild-type CrPV and CrPV-DcDV RNA levels were determined using 4.5 µL diluted cDNA for qPCR reactions with the SsoAdvanced Universal SYBR Green Supermix (BioRad, Hercules, California) in 10 µL reactions. Primers 19 and 20 were used for CrPV RNA and primers 21 and 22 were used for *D. citri* actin. All primers were used at a final concentration of 0.25 nM. The qPCR reaction conditions consisted of an initial denaturation at 98 °C for 2 min followed by 40 cycles at 95 °C for 10 s and 60 °C for 20 s. CrPV RNA levels were normalized based on the expression of *D. citri* actin using the 2^-ΔΔCt^ method (73). Technical triplicates were used for al qPCR reactions. Diluted cDNA from each day was pooled to assess retention of the insert in CrPV-DcDV as described above.

## Acknowledgements

We thank E. Matsumura for assistance and protocols for CrPV infections. We thank colleagues who provided us *D. citri* samples: W.O. Dawson, H. Wuriyanghan, H.-H Yeh, Y. Cen, A. M. Khan, M. A. Machado, T. da Silva, D. M. Galdeano, D. Jenkins, C. Higashi, A. Chow, D. Morgan, K. Pelz-Stelinski, K. Godfrey, B. Baker, G. Simmons, M. Keremane, and J. Buenahora. We thank S.W.-Ding for providing us with pCrPV-3. We also thank M-C. Saleh for critical reading of the manuscript. The U.S. Department of Agriculture provided funding to Bryce W. Falk under grant numbers 13-002NU-781 and 2015-70016-23011. This material is based upon work supported by the National Science Foundation Graduate Research Fellowship Program under Grant No. 1650042.

## Competing interests

The authors declare that they have no competing interests.

---

**Fig. S1.**
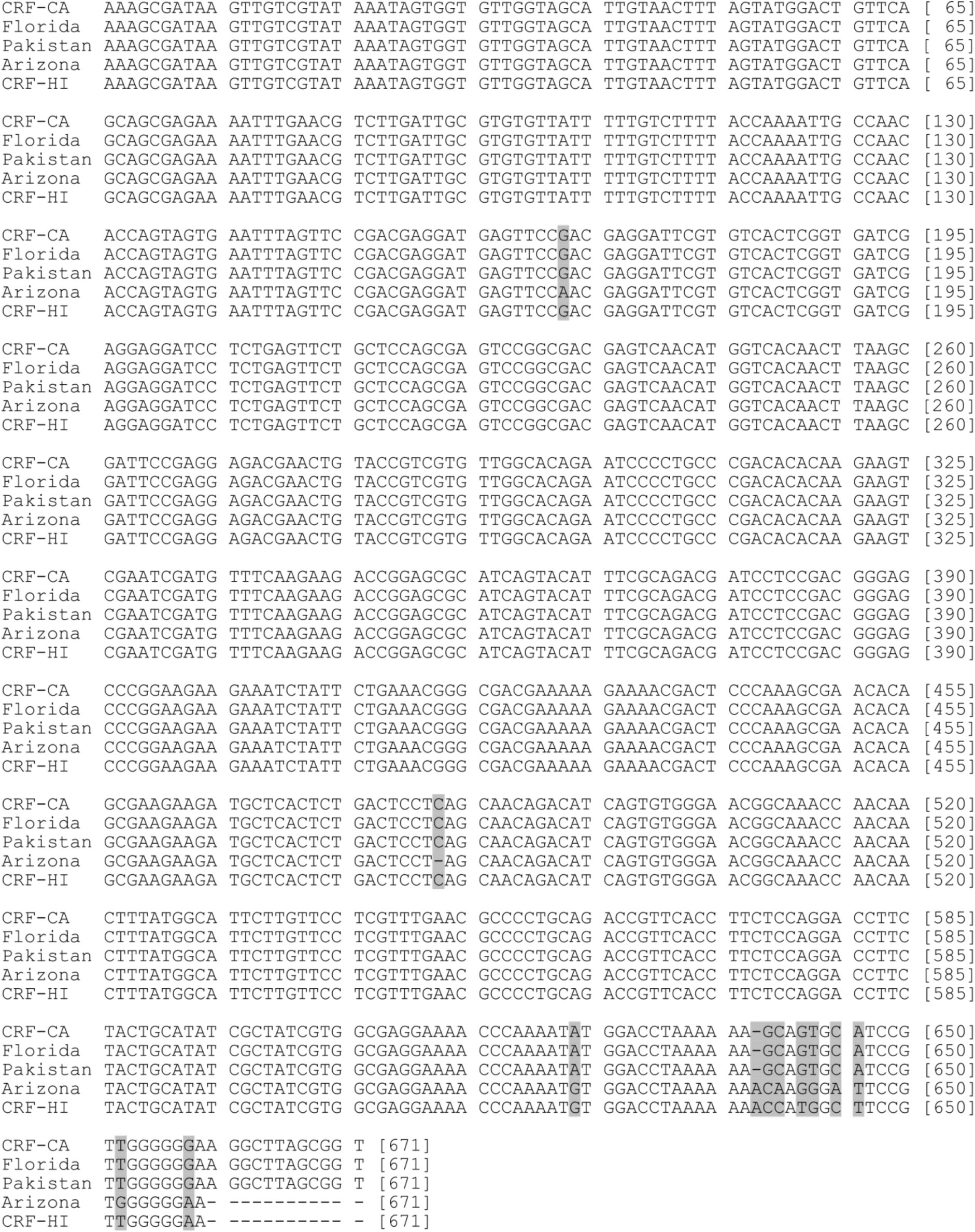
Alignment of the sequences of the ENS PCR products shown in Fig. 2. Sequences were aligned using ClustalW and mismatched nucleotides are shaded.

**Fig. S2.**
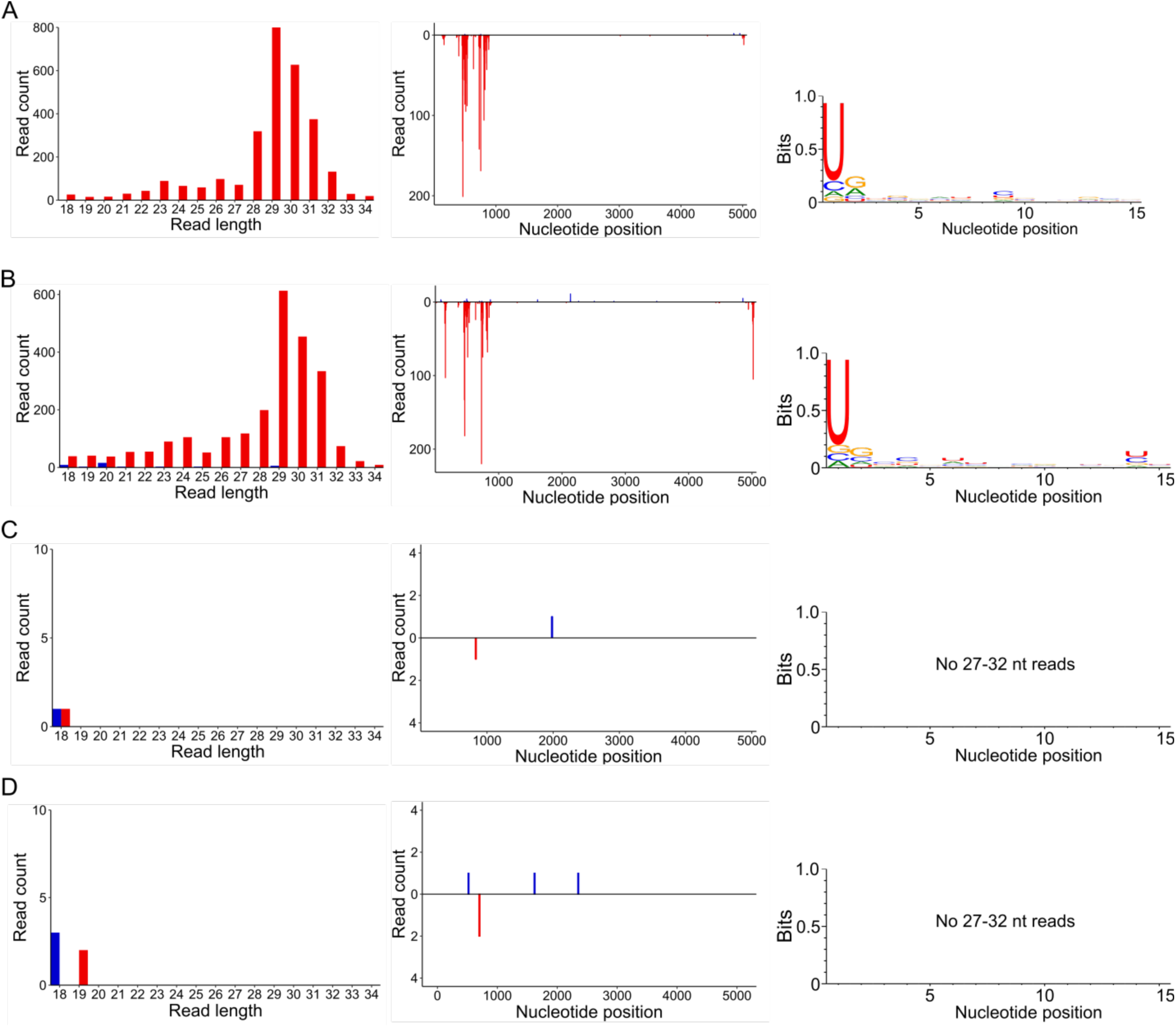
(A-D) Left: Length distribution of all sRNAs from various *D. citri* populations mapped to the DcDV genome. Red = antisense, blue = sense. Middle: Positions of all sRNAs from various *D. citri* populations mapped to the DcDV genome. sRNAs are shown as mapped to the genomic strand containing the coding sequence for the NS proteins. Red = antisense sRNAs, blue = sense sRNAs. Right: Sequence logo of 27-32 nt sRNAs shown in the left and middle panels. sRNAs were purified from field-collected insects from China (GenBank accession no. SRX1164134)(A), the US state of Florida (GenBank accession no. SRX1164135) (B), Brazil (GenBank accession no. SRX1164127)(C), or laboratory reared insects from CRF-Uru (D). Sequence data for (A-C) are described in (25, 26).

**Fig. S3.**
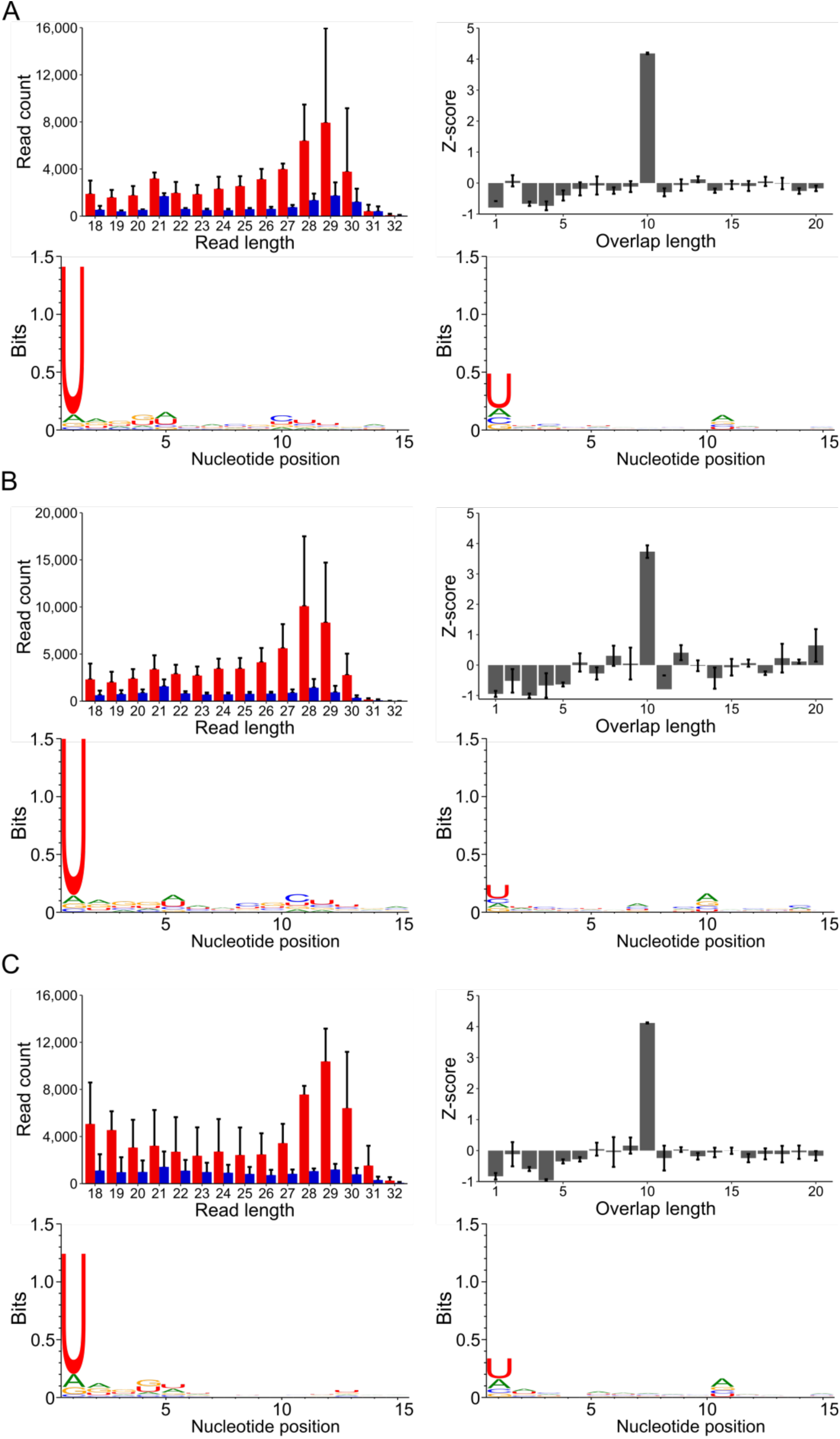
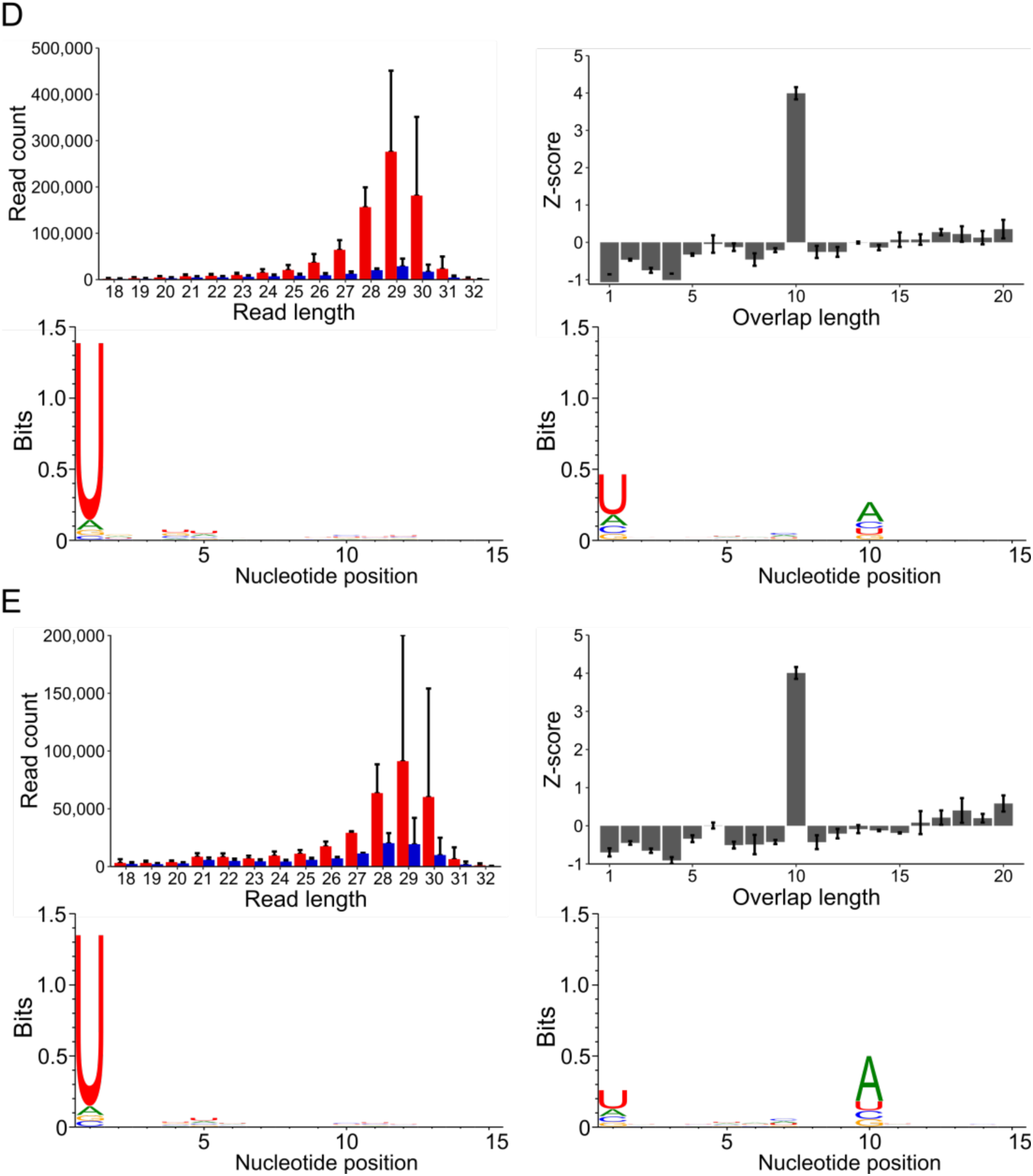
(A-E) Analysis of sRNAs from dissected CRF-CA *D. citri* tissues. Figures indicate mapping statistics for sRNAs mapping to all TEs identified in the *D. citri* genome. (A) sRNAs from three pools of 200 *D. citri* guts. (B) sRNAs from three pools of 200 *D. citri* heads. (C) sRNAs from three pools of hemolymph from 100 *D. citri* each. (D) sRNAs from three pools of 100 *D. citri* ovaries. (E) sRNAs from three pools of 100 *D. citri* testis. Upper left panels: length distribution of sRNAs mapping to TEs. Red = antisense, blue = sense. Read counts are an average of three independent libraries. Error bars indicate standard deviation. Upper right panels: Z-scores for the indicated overlap distances between the 5’ ends of complementary 27-32 nt sRNAs. Z-scores are an average of three independent libraries. Errors bars indicate the standard deviation. Lower panels: Sequence logos for 27-32 nt sRNAs mapping to TEs. Lower left = antisense, Lower right = sense. Data represents three pooled libraries.

**Fig. S4.**
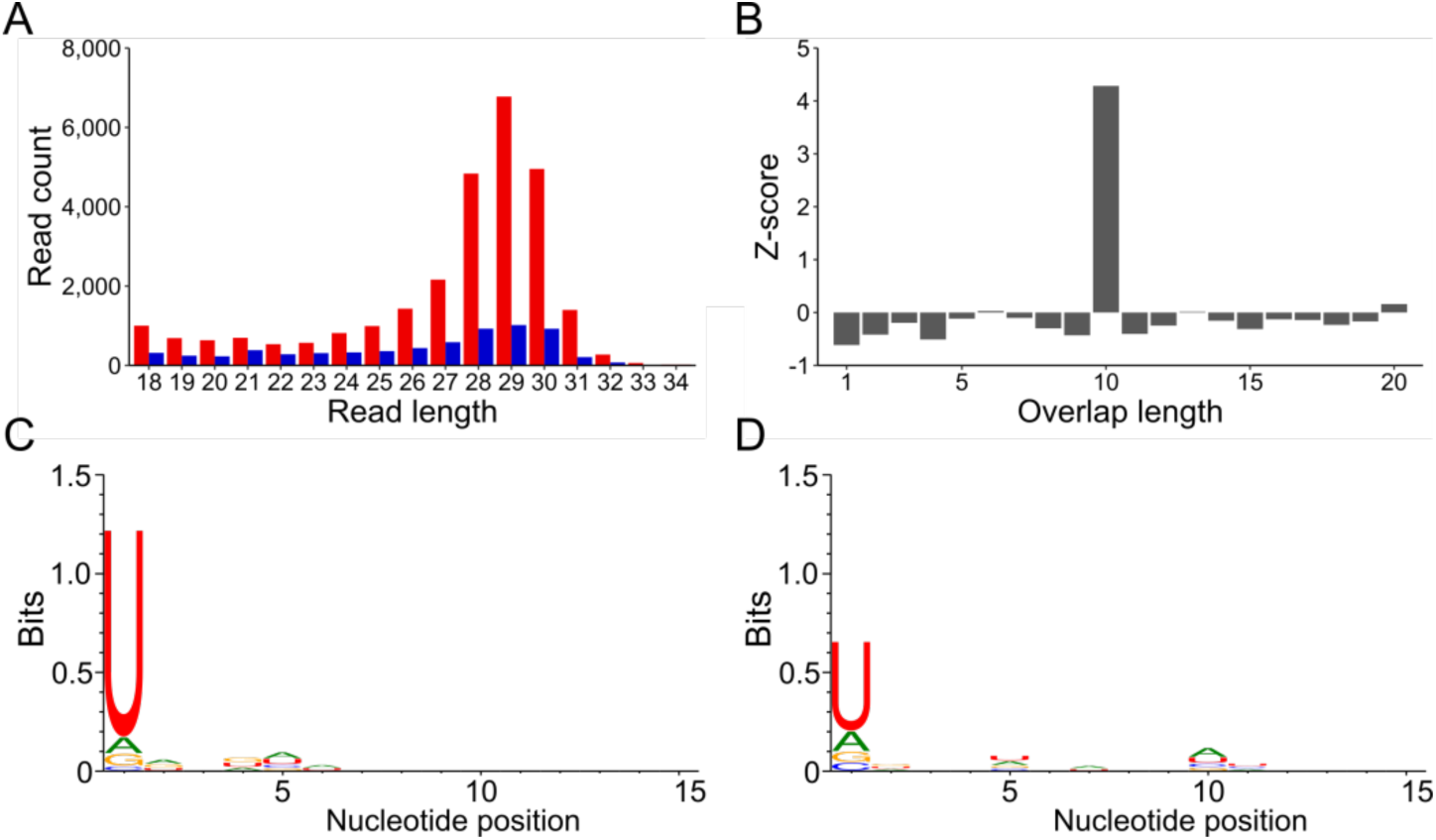
(A-E) Analysis of sRNAs from CRF-Uru *D. citri*. Figures indicate mapping statistics for sRNAs mapping to all TEs identified in the *D. citri* genome. (A) length distribution of sRNAs mapping to TEs. Red = antisense, blue = sense. (B) Z-scores for the indicated overlap distances between the 5’ ends of complementary 27-32 nt sRNAs. (C & D) Sequence logos for 27-32 nt sRNAs mapping to TEs. (C) Antisense sRNAs. (D) Sense sRNAs.

**Fig. S5.**
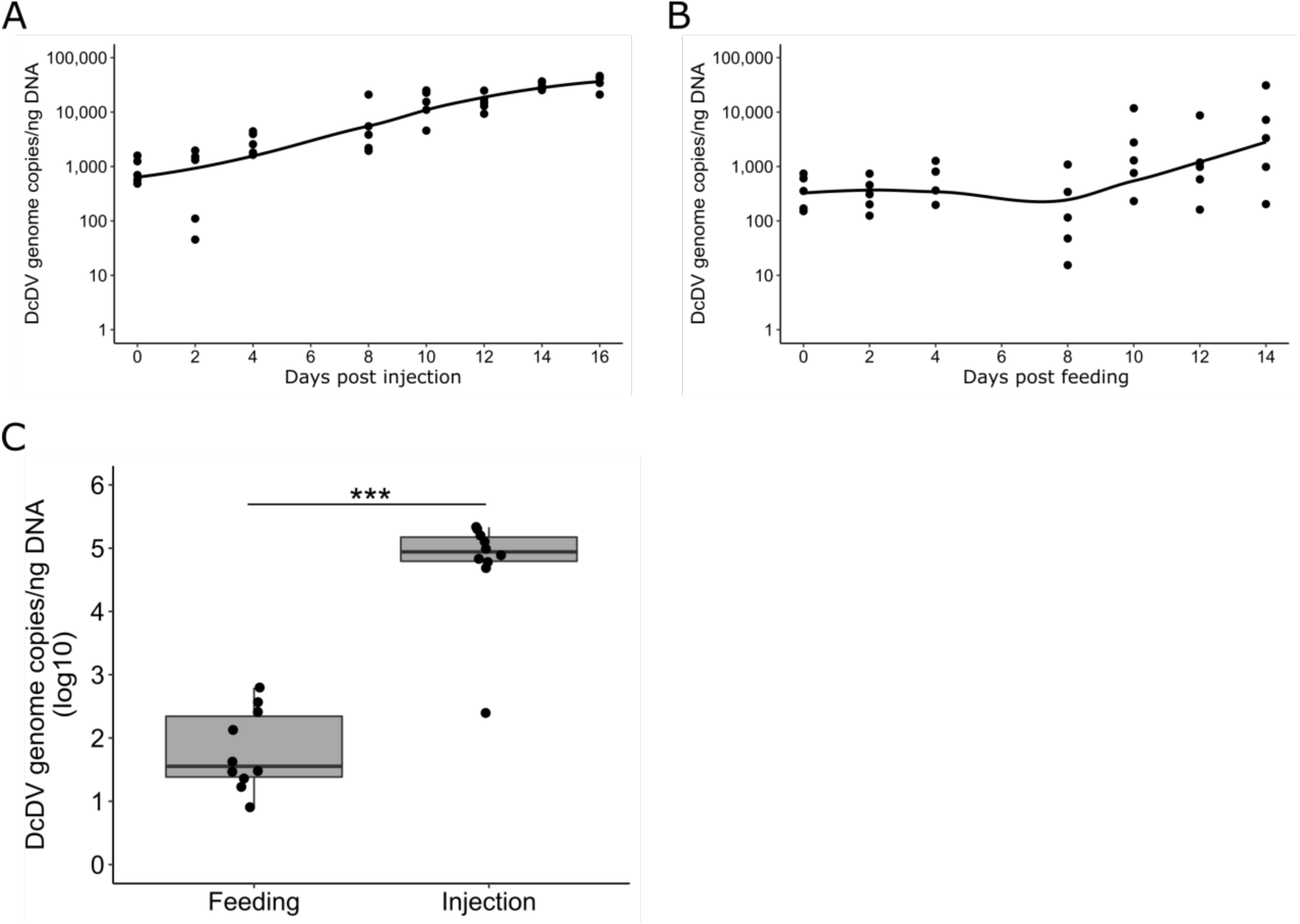
(A) Titer of DcDV following intrathoracic injection of virions into uninfected *D. citri* insects from CRF-Uru. Each data point represents three pooled insects (n=5). The curve was smoothed using loess regression. (B) Titer of DcDV following feeding of virions to uninfected *D. citri* insects from CRF-Uru. Virions present in a 15% sucrose artificial diet solution were acquired by *D. citri* for 96 hours by membrane feeding. After the feeding period, insects were moved to *Citrus macrophylla* plants and this represents day 0 post feeding. Each data point represents three pooled insects (n=5). The curve was smoothed using loess regression. (C) Titer of DcDV in F1 progeny of the insects shown in (A), designated by injection, and (B), designated by Feeding. n=10 individual insects. Individual insects were collected on the day of emergence to adulthood.

**Fig. S6.**
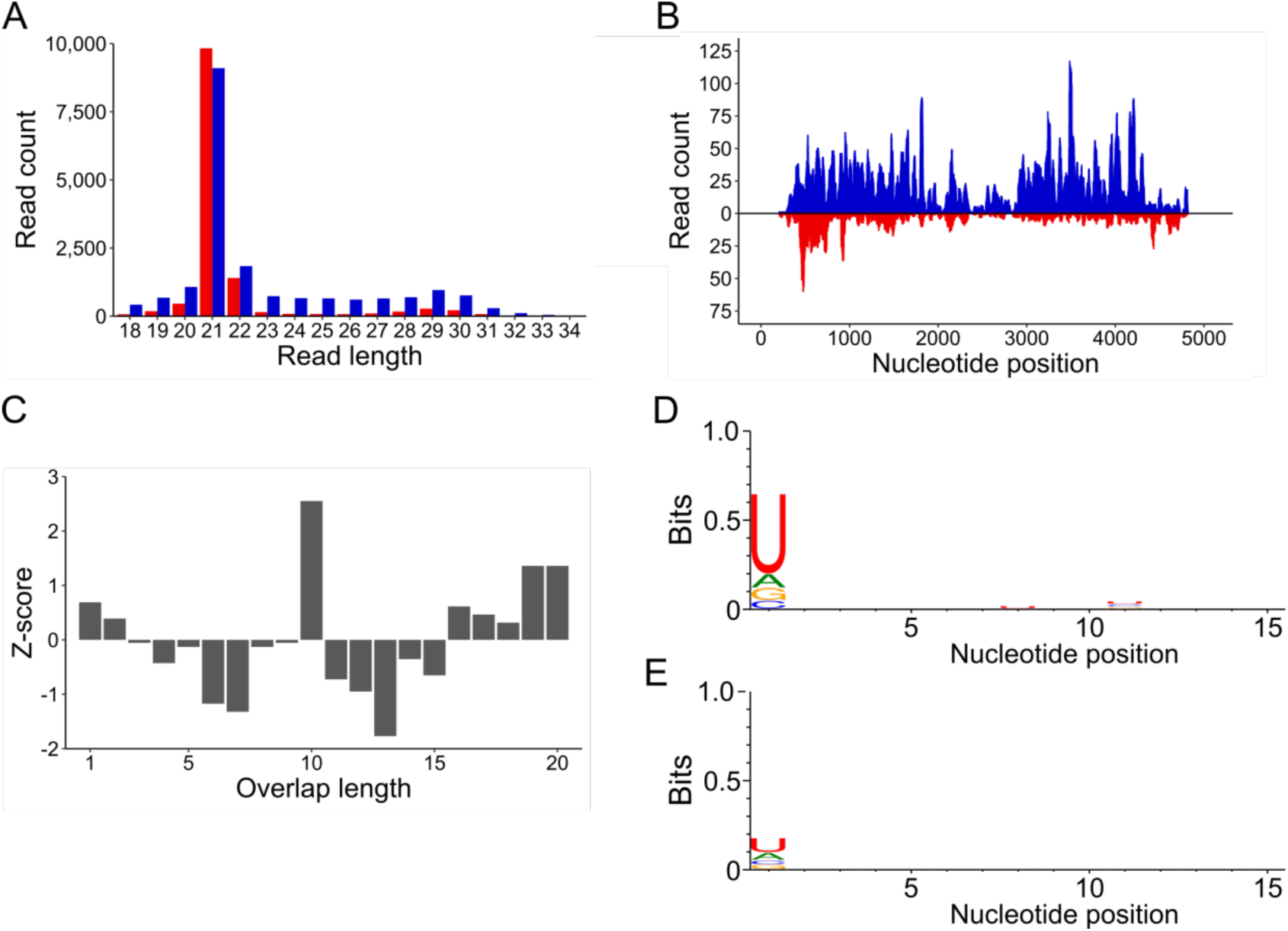
DcDV is targeted by ping-pong-dependent vpiRNAs in DcDV-infected *D. citri* insects from CRF-Uru. sRNAs were purified from the progeny of CRF-Uru *D. citri* that had been infected with DcDV by intrathoracic injection (see Fig. S5a & c). (A) Length distribution of sRNAs mapping to the DcDV genome in DcDV-infected *D. citri* insects from CRF-Uru. To account for the bidirectional transcription strategy of DcDV, sRNA mapping polarity was assigned from mapping location based on the start and stop positions of the canonical DcDV transcripts. Red = antisense sRNAs, blue = sense sRNAs. (B) Positions of 27-32 nt sRNAs shown in (A). Red = antisense sRNAs, blue = sense sRNAs. (C) Z-scores for the indicated overlap distances between the 5’ ends of complementary 27-32 nt sRNAs shown in (A). (D & E) Sequence logos for the 27-32 nt sRNAs shown in (A). Sequence logos for antisense sRNAs (D) or sense sRNAs (E) are shown.

**Fig. S7.**
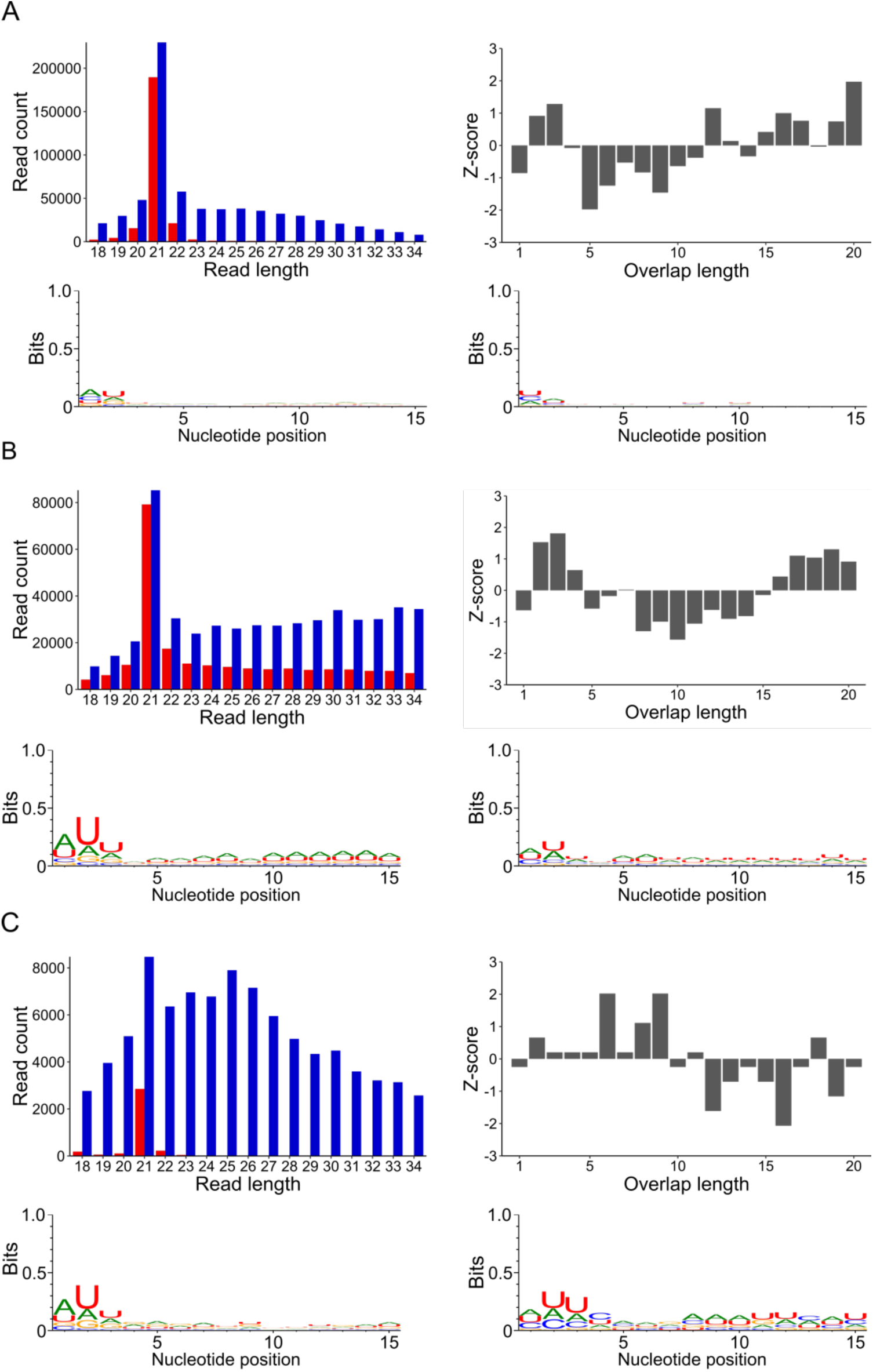
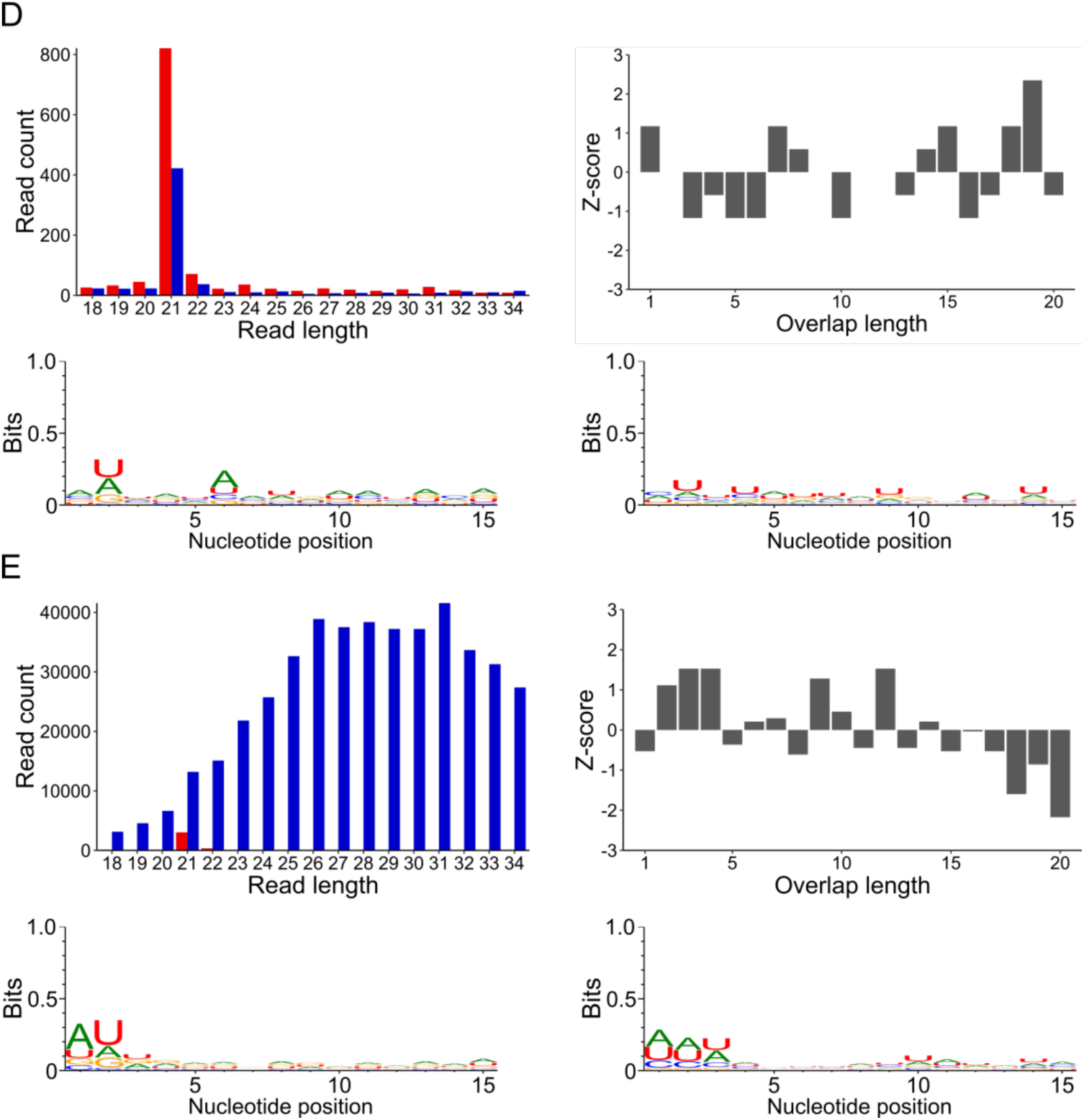
(A-E) Analysis of sRNAs from various field-collected *D. citri* populations infected with one or more *D. citri*-specific viruses. (A) *D. citri* from the US state of Florida infected with Diaphorina citri flavi-like virus (sRNA GenBank accession no. SRX1164135, viral genome GenBank accession no. NC_030453.1). (B) *D. citri* from China infected with Diaphorina citri reovirus (sRNA GenBank accession no. SRX1164134, viral genome GenBank accession no.’s KT698830.1, KT698831.1, KT698832.1, KT698833.1, KT698834.1, KT698836.1). (C) *D. citri* from China infected with Diaphorina citri picorna-like virus (sRNA GenBank accession no. SRX1164134, viral genome GenBank accession no. KT698837.1). (D) *D. citri* from China infected with Diaphorina citri bunyavirus (sRNA GenBank accession no. SRX1164134, viral genome GenBank accession no.’s KT698825.1, KT698824.1, KT698823.1). (E) *D. citri* from Brazil infected with Diaphorina citri picorna-like virus (sRNA GenBank accession no. SRX1164127, viral genome GenBank accession no. KT698837.1). Upper left panels: length distribution of sRNAs mapped to the various viral genomes. Red = antisense, blue = sense. Upper right panels: Z-scores for the indicated overlap distances between the 5’ ends of complementary 27-32 nt sRNAs mapping to opposite strands of the various viral genomes. Sequence data are described in (25, 26). Lower panels: Sequence logos for 27-32 nt sRNAs mapping to the various viral genomes. Lower left = antisense, Lower right = sense.

**Fig. 8.**
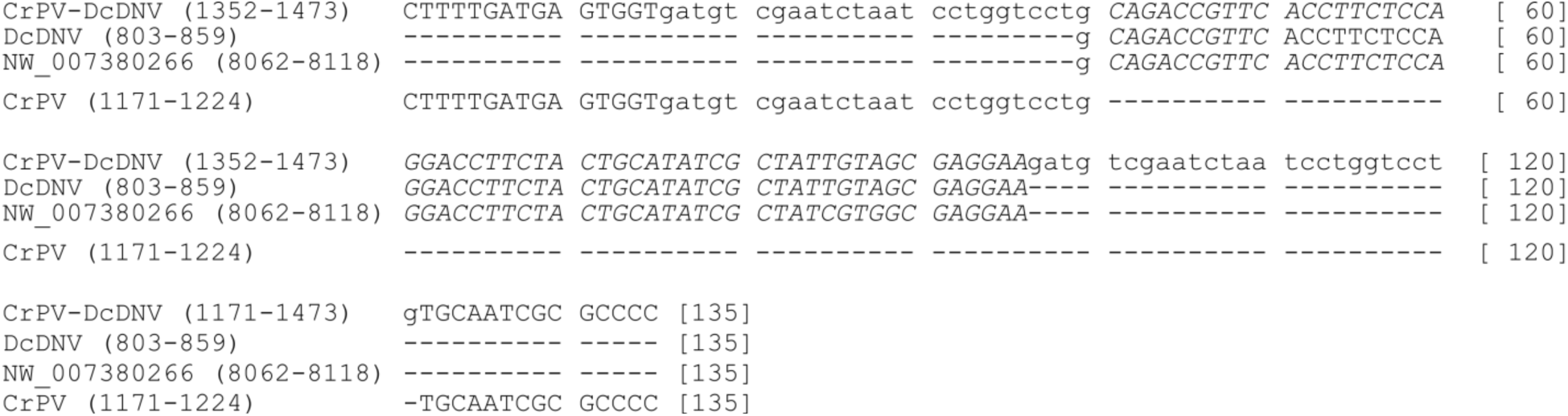
Alignment of a portion of the CrPV-DcDV genome, the recombinant DcDV sequence present within CrPV-DcDV, the region of ENS corresponding to the recombinant DcDV sequence present within CrPV-DcDV (represented by the GenBank accession no. of *D. citri* genomic scaffold 2850, NW_007380266), and a portion of the wild-type CrPV sequence. Numbers in parentheses indicate the nucleotide positions of the sequence shown. The CrPV 1A cleavage site is shown in lowercase letters. The recombinant DcDV sequence present within CrPV-DcDV and the corresponding region of ENS are shown in italics.

**Fig. S9.**
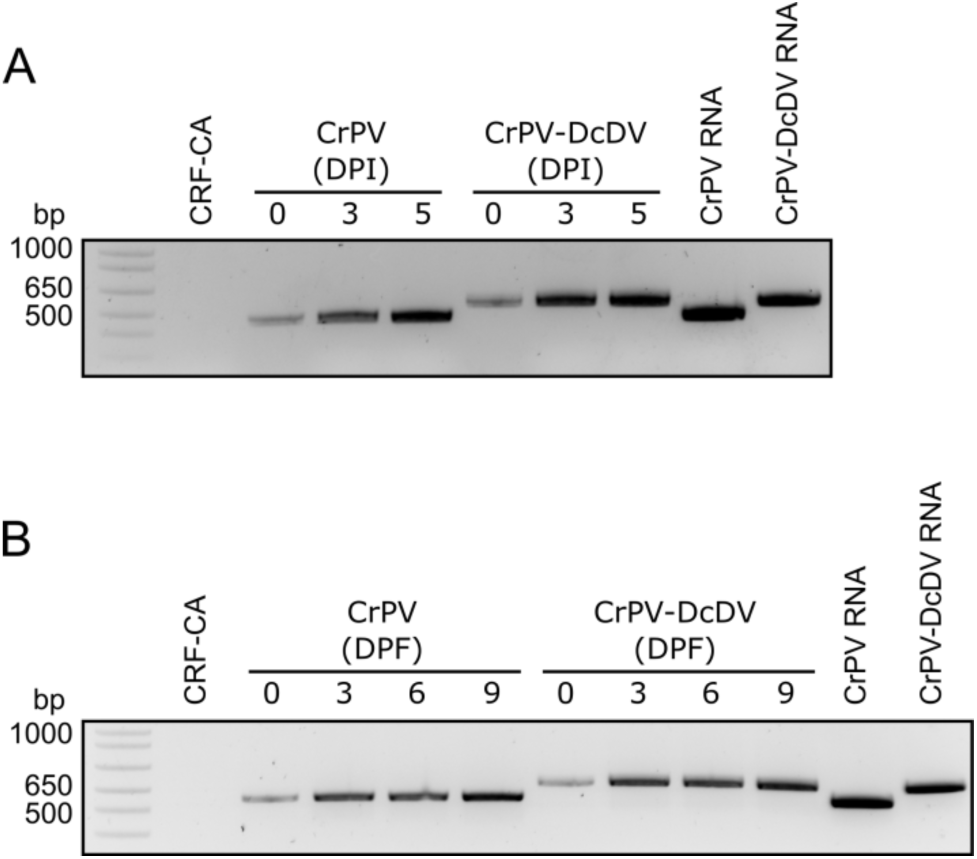
RT-PCR products produced using primers flanking the site into which recombinant DcDV sequence was inserted in CrPV-DcDV (primers 15 and 16). RNA from all biological replicates from each day shown in Fig. 4.6a (A) or Fig. 4.7a (B) was pooled and used as templates for RT-PCR. RNA from CRF-CA *D. citri* was used as a negative control. *In vitro* transcribed wild-type CrPV or CrPV-DcDV RNA was used as a positive control. DPI = days post injection. DPF = days post feeding.

**Fig. S10.**
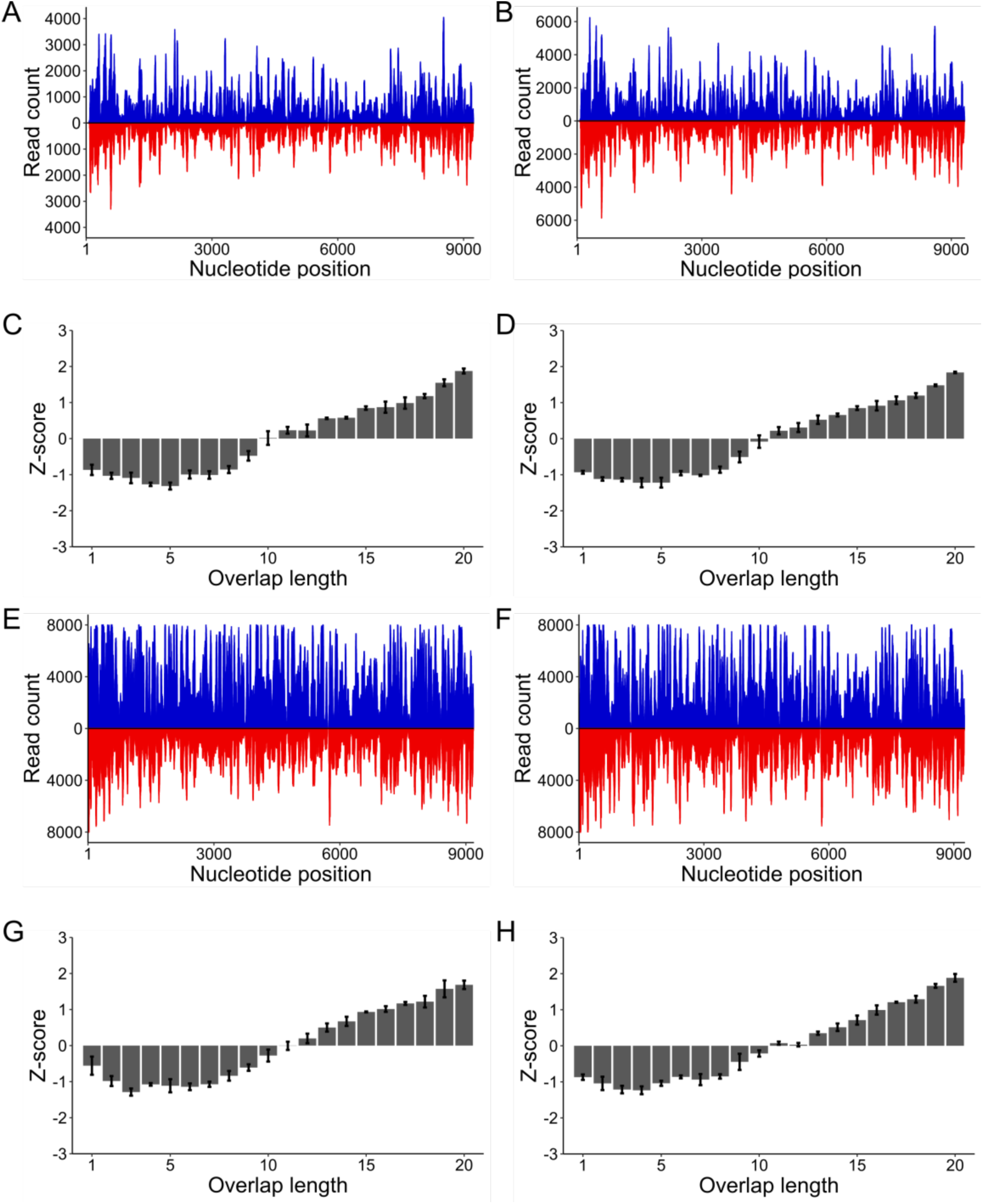
(A & B) Positions of 21 nt sRNAs mapping to the wild-type CrPV genome (A) or the CrPV-DcDV genome (B) during infection of CRF-CA *D. citri* initiated by intrathoracic injection of virions. Red = antisense sRNAs, blue = sense sRNAs. Reads counts are an average of three independent libraries. (C & D) Z-scores for the indicated overlap distances between the 5’ ends of complementary 27-32 nt sRNAs mapping to opposite strands of the wild-type CrPV genome (C) or the non-recombinant portion of the CrPV-DcDV genome (D) during infection of CRF-CA *D. citri* initiated by intrathoracic injection of virions. Z-scores represent the average of three independent libraries. Error bars indicate standard deviation. (E & F) Positions of 21 nt sRNAs mapping to the wild-type CrPV genome (E) or the CrPV-DcDV genome (F) during infection of CRF-CA *D. citri* initiated by oral acquisition of virions. Red = antisense sRNAs, blue = sense sRNAs. Reads counts are an average of three independent libraries. (G & H) Z-scores for the indicated overlap distances between the 5’ ends of complementary 27-32 nt sRNAs mapping to opposite strands of the wild-type CrPV genome (G) or the non-recombinant portion of the CrPV-DcDV genome (H) during infection of CRF-CA *D. citri* initiated by oral acquisition of virions. Z-scores represent the average of three independent libraries. Error bars indicate standard deviation.

**Table S1.**
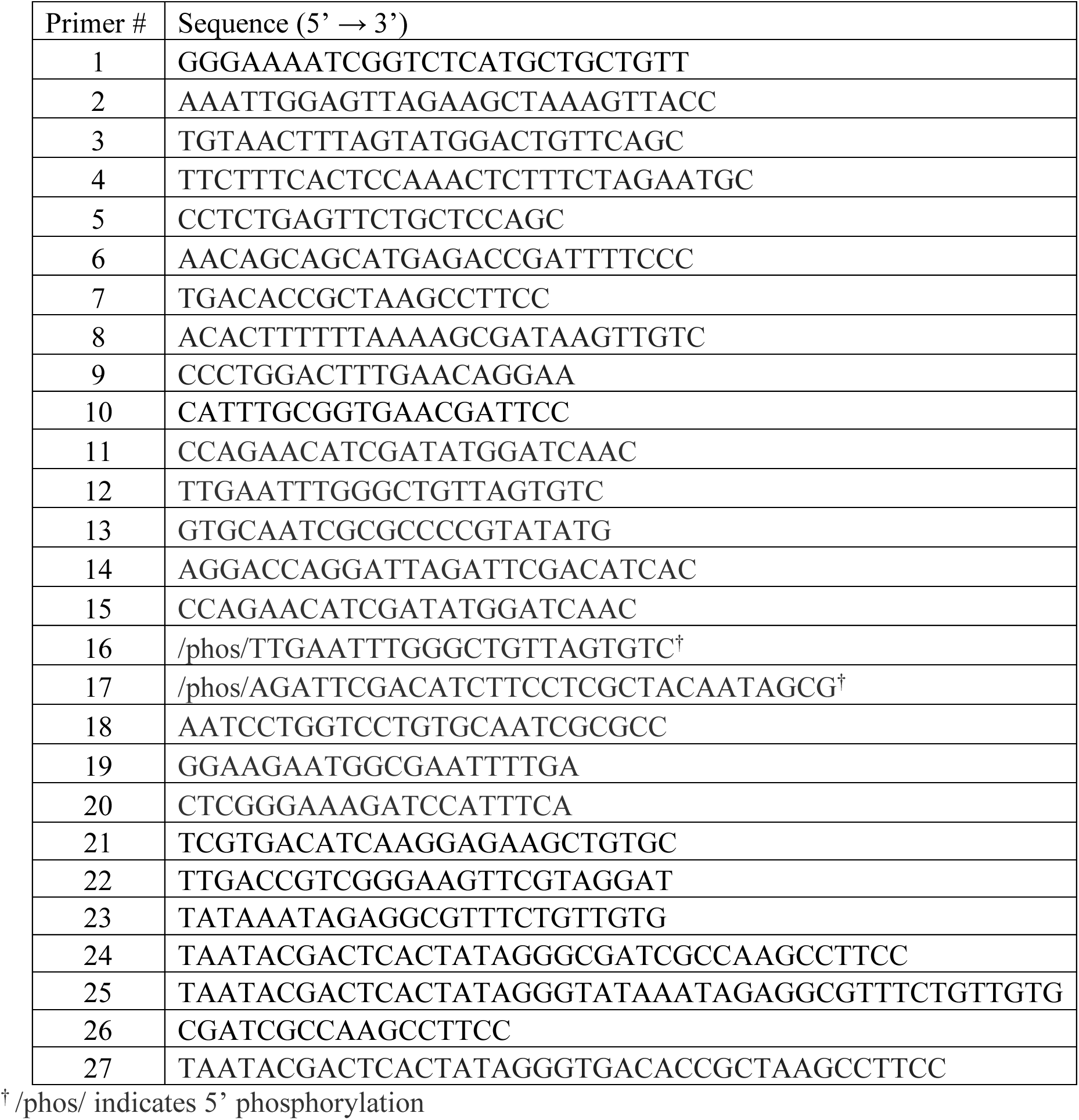

## Notes

### Competing Interest Statement

The authors have declared no competing interest.

